# Concomitant suppression of COX-1 and COX-2 is insufficient to induce enteropathy associated with chronic NSAID use

**DOI:** 10.1101/2024.11.22.624882

**Authors:** Kayla Barekat, Soumita Ghosh, Christin Herrmann, Karl Keat, Charles-Antoine Assenmacher, Ceylan Tanes, Naomi Wilson, Arjun Sengupta, Ujjalkumar Subhash Das, Robin Joshi, Marylyn D. Ritchie, Kyle Bittinger, Aalim Weljie, Ken Cadwell, Frederic D. Bushman, Gary D. Wu, Garret A. FitzGerald, Emanuela Ricciotti

## Abstract

Nonsteroidal anti-inflammatory drugs (NSAIDs) are the most widely used medications for the management of chronic pain; however, they are associated with numerous gastrointestinal (GI) adverse events. Although many mechanisms have been suggested, NSAID-induced enteropathy has been thought to be primarily due to inhibition of both cyclooxygenases (COX) -1 and -2, which results in suppression of prostaglandin synthesis. Yet surprisingly, we found that concomitant postnatal deletion of *Cox-1* and *-2* over 10 months failed to cause intestinal injury in mice unless they were treated with naproxen or its structural analog, phenylpropionic acid, which is not a COX inhibitor. *Cox* double knockout mice exhibit a distinct gut microbiome composition and cohousing them with controls rescues their dysbiosis and delays the onset of NSAID-induced GI bleeding. In both the UK Biobank and All of Us human cohorts, coadministration of antibiotics with NSAIDs is associated with an increased frequency of GI bleeding. These results show that prostaglandin suppression plays a trivial role in NSAID-induced enteropathy. However, *Cox* deletion causes dysbiosis of the gut microbiome that amplifies the enteropathic response to NSAIDs.

## Introduction

Nonsteroidal anti-inflammatory drugs (NSAIDs) are the most widely used medications for the management of chronic pain and inflammation, with tens of millions of people worldwide consuming them daily.^[1, 2]^ NSAIDs are not addictive like opiates, but they are still associated with gastrointestinal (GI)^[3]^ and cardiovascular (CV)^[4]^ adverse events. Endoscopic studies have estimated the prevalence of inflammation in 60-70% of chronic users, blood loss and anemia in 30%, and mucosal ulceration in 30-40%.^[5, 6]^ Notably, there is a wide range of inter-individual variation in response to NSAIDs – in both safety and efficacy – which cannot be fully explained by genetic differences in host metabolizing enzymes.^[7, 8]^ While the mechanisms underlying NSAID-induced gastropathy are well-established,^[9–17]^ NSAID-induced enteropathy remains poorly understood, and there is still no treatment or preventative solution for the intestinal symptoms.

The anti-inflammatory, analgesic efficacy of NSAIDs is attributable to suppression of prostaglandins by cyclooxygenases (COX) -1 and -2.^[18, 19]^ COX-1 is the more constitutively expressed isoform while COX-2 is induced by inflammatory and mitogenic stimuli,^[20]^ albeit that these distinctions are not absolute – COX-1 can be up-regulated under certain circumstances,^[21–24]^ and COX-2 is detectable in some tissues unchallenged.^[25–29]^ Enteropathy has been attributed to inhibition of COX-1-dependent synthesis of prostaglandin (PG) E_2_ and prostacyclin (PGI_2_) in intestinal epithelial cells, which are involved in protection of the mucosal barrier.^[30]^ This is compounded by inhibition of COX-1-dependent thromboxane (Tx) A_2_ formation in platelets, predisposing to hemorrhage.^[31]^ However, pharmacological inhibition or genetic deletion of COX-1 alone does not result in spontaneous enteric lesions.^[32, 33]^ Conventional deletion of both COXs is embryonically lethal, but combined genetic deletion of one and pharmacological inhibition of the other (as well as pharmacological inhibition of both) results in enterotoxicity.^[33–35]^ COX-2-derived PGs also play key roles in intestinal mucosal defense^[36]^ and injury repair.^[37, 38]^ Yet similar to the case of COX-1 suppression alone, COX-2 suppression alone results in little to no enteric injury relative to nonselective NSAIDs in mice.^[39–42]^ In humans, clinical trials in arthritis have shown that selective inhibition of COX-2 roughly halves the incidence of GI bleeds compared to mixed inhibitors,^[43, 44]^ and although expression of both enzymes is upregulated in inflammatory synovia,^[45]^ comparative efficacy trials of selective vs nonselective NSAIDs have not been performed.

While prostaglandin suppression has been thought to be the dominant cause of NSAID enteropathy, additional factors including direct chemical toxicity,^[46–48]^ disruption of mitochondrial function,^[49, 50]^ intestinal dysbiosis,^[51–57]^ and bile acid toxicity^[58–60]^ have also been implicated. To address further the mechanisms that underlie NSAID enteropathy, we developed two reagents: 1) mice in which deletion of both enzymes is achieved postnatally, thereby circumventing the critical role of COXs in pregnancy and embryonic development;^[61, 62]^ and 2) a chronic dosing model with the NSAID naproxen, where drug exposure and prostaglandin suppression corresponds to what is seen in humans. To our surprise there was minimal evidence of enteropathy consequent to *Cox* deletion, whereas naproxen-induced GI toxicity was amplified in the double knockouts (DKOs). As prostaglandin suppression was comparable between the DKOs and naproxen treatment, it is not the explanation for the NSAID enterotoxicity. Rather, drug interactions with the gut microbiome are of more importance.

## Results

We developed a tamoxifen-inducible universal Cre-recombinase mouse line with loxP sites flanking both *Cox-1* and *Cox-2*, respectively. We first confirmed that these *Cox-1^fx/fx^ Cox-2^fx/fx^ CMV-Cre^+/-^* (“*Cox-* DKO”) mice exhibited reduced expression of the genes in two separate tissues (Supplemental Figure 1A). We next confirmed prostaglandin suppression *in vivo* in the DKOs by measuring urinary metabolites over a sustained period (Figure 1A). This difference from *Cre^-/-^* control littermates was evoked further by stimulation of prostaglandin synthesis with lipopolysaccharide (LPS), which induces *Cox* expression (Figure 1B). Blinded assessment of tissue histology (Table 1, Supplemental Figure 2) failed to detect gastrointestinal ulcers (Figure 1C) or visible signs of enteropathy (Figure 1D) ten months post-tamoxifen induced genetic deletion (approximately one year of age) in the DKOs. Furthermore, no blood was detected in the stool of any individual mouse with weekly hemoccult testing (Figure 1E). Thus, marked suppression of prostaglandin synthesis alone does not result in gastroenteropathy.

**Figure 1:**
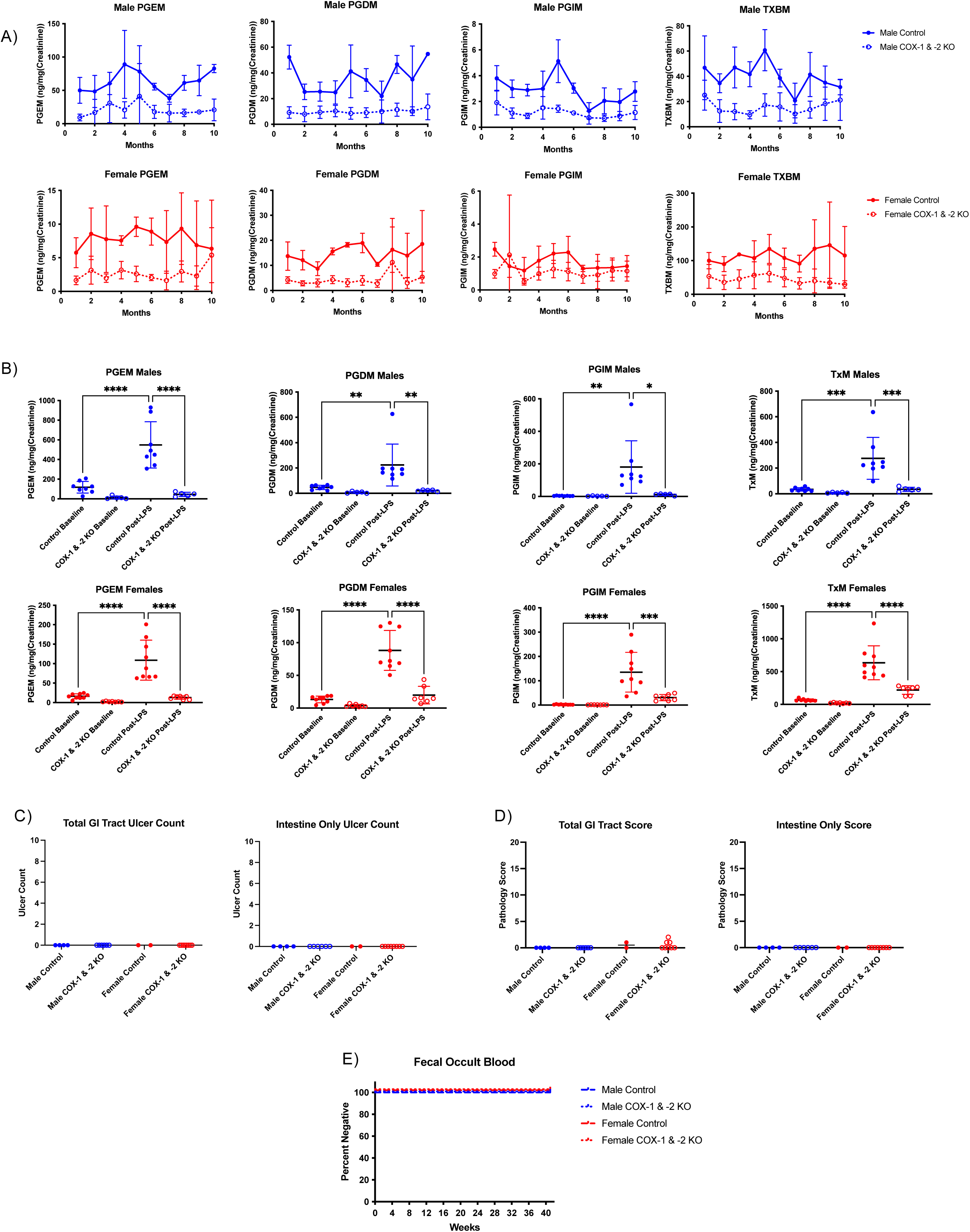
Postnatal deletion of *Cox-1* and *-2* reveals that chronic prostaglandin depletion alone is not sufficient to induce spontaneous GI injury in mice. (**A**) Longitudinal tracking of universal inducible *Cox*-DKO mice was conducted over 10 months post-tamoxifen exposure (administered at 10-12 weeks of age). Monthly urinary prostaglandin metabolite levels measured by LC-MS/MS, analyzed separately for each sex. n = 14 *Cox*-DKO mice and 6 *Cre^-/-^* controls. (**B**) Attempted induction of prostaglandin synthesis in *Cox*-DKO mice via lipopolysaccharide (LPS) challenge. Mice received 1 mg/kg body weight LPS by IP injection, and urine was collected overnight. n = 7-9 mice per group. Urinary prostaglandin metabolites measured by LC-MS/MS for both male and female *Cox*-DKO mice and *Cre^-/-^* controls at baseline vs post-LPS exposure. *p<0.05, **p<0.01, ***p<0.001, ****p<0.0001 by one-way ANOVA. (**C**) Total ulcer count throughout the entire GI tract (including stomach, small intestine, and large intestine) for longitudinal tracking experiment, as counted by a blinded pathologist. Each data point represents a single mouse. (**D**) Qualitative / semi-quantitative pathology scores throughout the entire GI tract for longitudinal tracking experiment. Each data point represents a single mouse. (**E**) Weekly hemoccult test results plotted as a Kaplan-Meier curve for percentage of each group that tested negative for blood in the stool. Any individual that tested positive would be marked positive for that first week and all subsequent weeks, resembling a survival curve.

**Table 1:** Gastrointestinal pathology scoring criteria. Qualitative / semi-quantitative scoring system for assessment of gastrointestinal pathology.

|  |  |  | Comment |
| --- | --- | --- | --- |
| Ulcer Distribution |  |  |  |
|  | None | 0 |  |
|  | Focal | 1 |  |
|  | Multifocal | 2 |  |
| Numbers of Ulcers |  |  | Combined number of ulcers visualized in the sections examined |
|  | Numbers Visualized | 0 - 5 |  |
| Ulcer Severity |  |  | Based on the most severe lesions in each individual section |
|  | Superficial | 1 |  |
|  | Up to Muscularis Mucosa | 2 |  |
|  | Transmural | 3 |  |
| Peritonitis/Serositis |  |  |  |
|  | No | 0 |  |
|  | Yes | 1 |  |
| Inflammation |  |  | Based on the most severe lesions |
|  | None | 0 |  |
|  | Minimal | 1 | Scattered inflammatory cells near the ulcer(s) or in the lamina propria |
|  | Mild | 2 | Few inflammatory cells near the ulcer(s) or in the lamina propria |
|  | Moderate | 3 | Moderate numbers of inflammatory cells near the ulcer(s) or in the lamina propria, occasionally forming sheets and aggregates |
|  | Severe | 4 | Large numbers of inflammatory cells near the ulcer(s) or in the lamina propria, occasionally forming sheets and aggregates |
| Total Score |  |  | Sum of all individual scores |

Naproxen is a nonselective NSAID that can elicit GI injury in both mice and humans while celecoxib, selective for inhibition of COX-2, is associated with a reduced GI toxicity profile.^[44]^ C57BL/6 wildtype mice were fed a custom diet with either naproxen (230 mg/kg), celecoxib (100 mg/kg), or no drug (control) dry-mixed into it and were allowed to feed *ad libitum* for 3 full weeks prior to tissue collection. Plasma drug concentrations at steady state for both NSAIDs correspond to human systemic drug exposure under chronic dosing conditions (Supplemental Figure 3A).

After 3 weeks exposure, naproxen (but not celecoxib) caused GI ulcers and inflammation in some mice, mostly in the small intestine (Figure 2A). Some of the mice with enteropathy showed signs of anemia (Figure 2B), neutrophilia and lymphopenia (Figure 2C), an increase in spleen weight relative to body weight (Figure 2D), and decreases in albumin, alkaline phosphatase, and total serum proteins (Figure 2E). In summary, levels of systemic naproxen exposure that correspond to those attained clinically result in ulceration, inflammation, GI bleeding, and protein deficiency characteristic of NSAID-induced enteropathy in humans.

**Figure 2:**
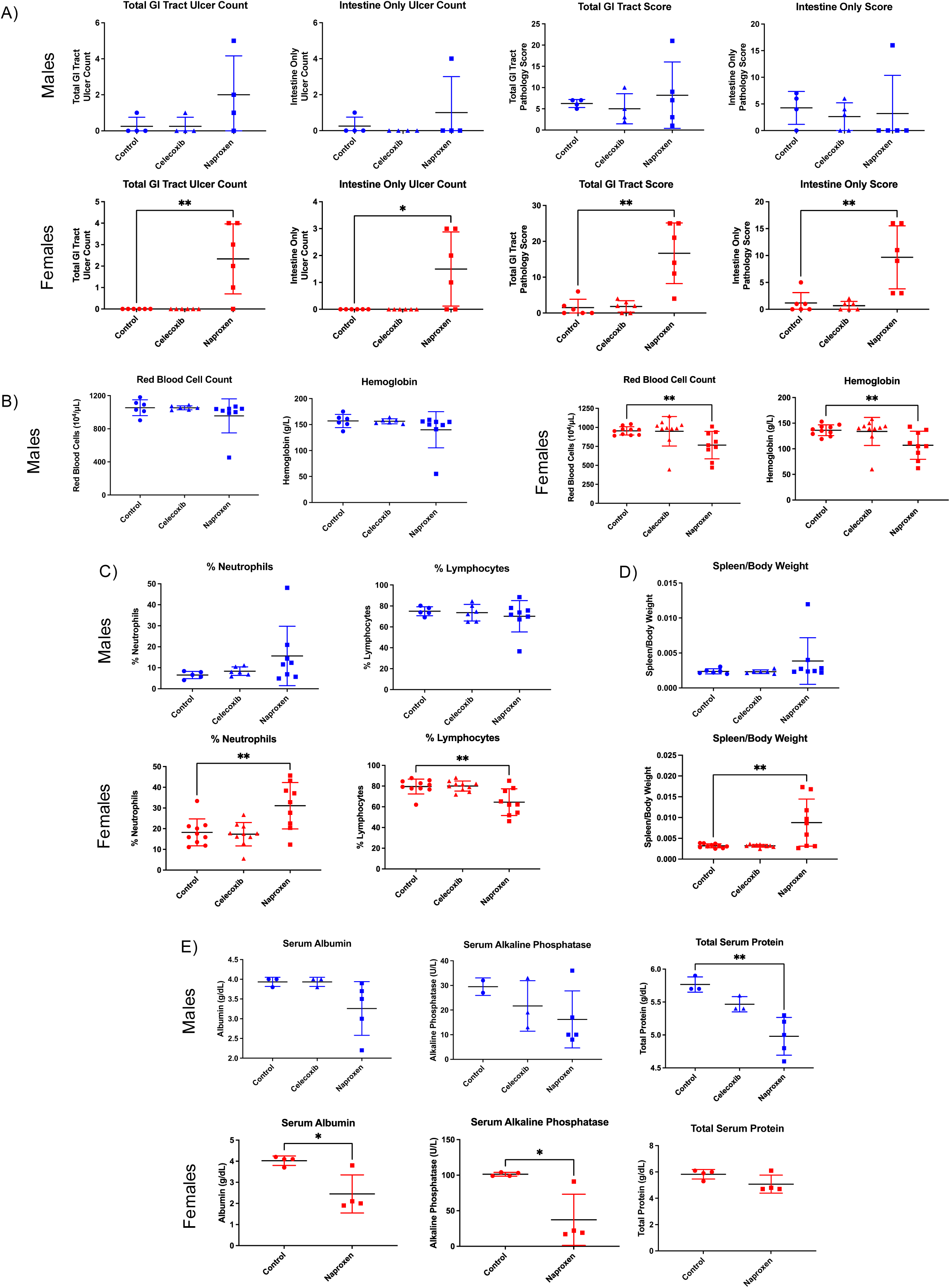
Chronic naproxen dosing in mice mimics NSAID enteropathy in humans. Wildtype C57BL/6J mice were treated with either control diet, celecoxib diet (100 mg/kg), or naproxen diet (230 mg/kg) and allowed to feed *ad libitum* for 3 weeks prior to tissue collection. (**A**) Ulcer count for total GI tract vs small intestine only, as well as pathology score for total GI tract vs small intestine only; males and females graphed separately. n = 4-6 mice per group. *p<0.05, **p<0.01 by unpaired t test. (**B**) Red blood cell count and hemoglobin measured by complete blood count (CBC); males and females graphed separately. n = 6-9 mice per group. **p<0.01 by unpaired t test. (**C**) Percentage of neutrophils and lymphocytes relative to all white blood cells measured by CBC; males and females graphed separately. N = 6-9 mice per group. **p<0.01 by unpaired t test. (**D**) Spleen weight relative to total body weight; males and females graphed separately. n = 6-9 mice per group. **p<0.01 by unpaired t test. (**E**) Serum proteins and liver enzymes measured by clinical automated chemistry analyzer; males and females graphed separately. n = 3-5 mice per group. *p<0.05, **p<0.01 by unpaired t test.

Urinary PGE-M and PGD-M were suppressed in both sexes both by *Cox* deletion and naproxen treatment, while urinary Tx-M was depressed in both sexes in the DKOs but not on naproxen. Addition of naproxen to the DKO mice did not significantly further depress prostaglandin synthesis (Figure 3A). DKO mice exhibited blood in the stool (Figure 3B), weight loss (Figure 3C), and ulcers (Figure 3D) only when they were treated for 3 weeks with naproxen. However, naproxen-induced GI injury was amplified when given on a background of *Cox* depletion. This was also true of indirect indices of GI blood loss – reticulocytosis and anemia (Figure 3E), neutrophilia and lymphopenia (Figure 3F), and increased spleen weight (Figure 3G).

**Figure 3:**
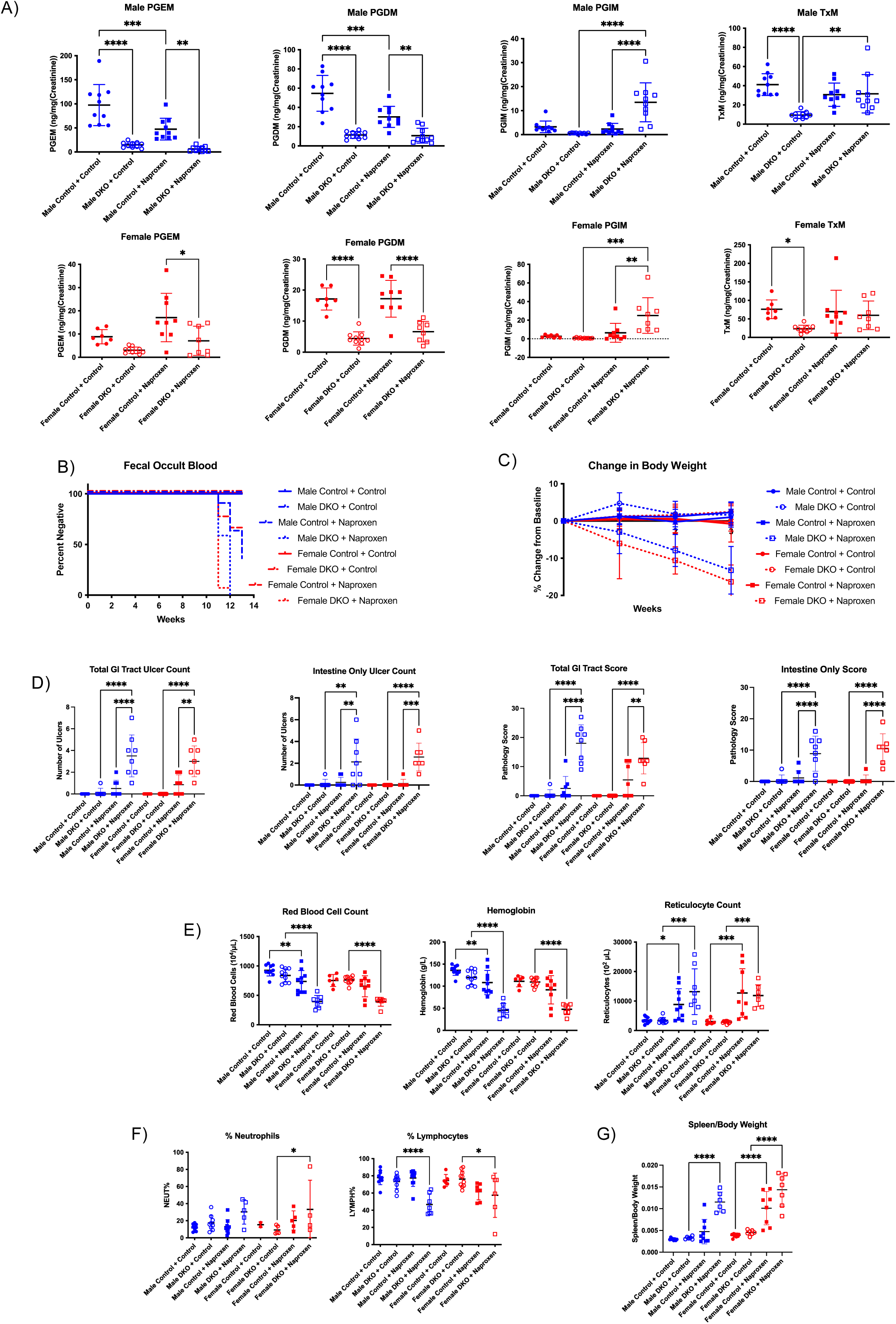
Naproxen enteropathy is exacerbated in *Cox*-DKO mice. *Cox*-DKO and *Cre^-/-^* control mice were treated with either control diet or naproxen diet (230 mg/kg) and allowed to feed *ad libitum* for 3 weeks prior to tissue collection. n = 7-10 mice per group for entire figure. (**A**) Urinary prostaglandin metabolites measured by LC-MS/MS, analyzed separately for each sex. *p<0.05, **p<0.01, ***p<0.001, ****p<0.0001 by one-way ANOVA. (**B**) Weekly hemoccult test results plotted as a Kaplan-Meier curve for percentage of each group that tested negative for blood in the stool. Any individual that tested positive would be marked positive for that first week and all subsequent weeks, resembling a survival curve. (**C**) Percent change in body weight relative to baseline body weight. (**D**) Ulcer count for total GI tract vs small intestine only, as well as pathology score for total GI tract vs small intestine only; males and females graphed separately. **p<0.01, ***p<0.001, ****p<0.0001 by one-way ANOVA. (**E**) Red blood cells, hemoglobin, and reticulocytes measured by CBC. *p<0.05, **p<0.01, ***p<0.001, ****p<0.0001 by one-way ANOVA. (**F**) Percentage of neutrophils and lymphocytes relative to all white blood cells measured by CBC. *p<0.05, ****p<0.0001 by one-way ANOVA. (**G**) Spleen weight relative to total body weight. ****p<0.0001 by one-way ANOVA.

As an additional control, we repeated the chronic naproxen dosing experiment using R-2-phenylpropionic acid (PPA), which is a weakly acidic structural analog of naproxen that lacks its COX inhibitory property.^[63]^ The same concentration of PPA (230 mg/kg) as naproxen was dry mixed into the diet and mice were allowed to feed *ad libitum* for 3 weeks prior to tissue harvest. As expected, PPA did not depress prostaglandin synthesis (Supplemental Figure 4A) but did cause anemia (Supplemental Figure 4B) and enteropathy (Supplemental Figure 4C), again exacerbated by *Cox* depletion as observed with naproxen. Unlike naproxen, PPA did not cause weight loss (Supplemental Figure 4D). Many of the *Cox*-DKOs on PPA exhibited some degree of injury in both the small intestine and stomach, whereas the controls were mostly unaffected. In summary, although prostaglandin suppression alone – by *Cox* depletion – does not cause enteropathy, it enhances the sensitivity of the mice to enteropathy induced by naproxen or its structural analog PPA.

Given the recognized involvement of the gut microbiome in NSAID-associated enteropathy, ^[51, 53, 54, 64, 65]^ we performed 16S rRNA sequencing of feces from the DKOs and controls. Littermates were separated by genotype shortly after weaning, prior to tamoxifen exposure. The heatmap summary shows that the baseline gut microbiome differed by genotype (Figure 4A), with the *Cox*-DKOs displaying an increased relative abundance of *Prevotella* and a decreased relative abundance of *Turicibacter* and *Dubosiella*. Both sexes exhibited distinct microbiome composition by genotype taking abundance of taxa into account (weighted UniFrac distance, male p=0.001, female p=0.005), but only females showed differences based on presence vs absence of taxa (unweighted UniFrac distance, male p=0.08, female p=0.02) (Figure 4B). After naproxen treatment, microbiome composition still differed by genotype for both the weighted and unweighted UniFrac distance metrics (Figure 4C, male p=0.001, female p=0.005 for weighted UniFrac; and male p=0.03, female p=0.03 for unweighted UniFrac). Microbiome differences did not manifest with treatment alone, and the effect of genotype was independent of treatment. The Firmicutes:Bacteroidota ratio was also altered in the *Cox*-DKO mice (Figure 4D, baseline: male p=0.004, female p=0.03; post-treatment: male p=0.004, female p=0.01).

**Figure 4:**
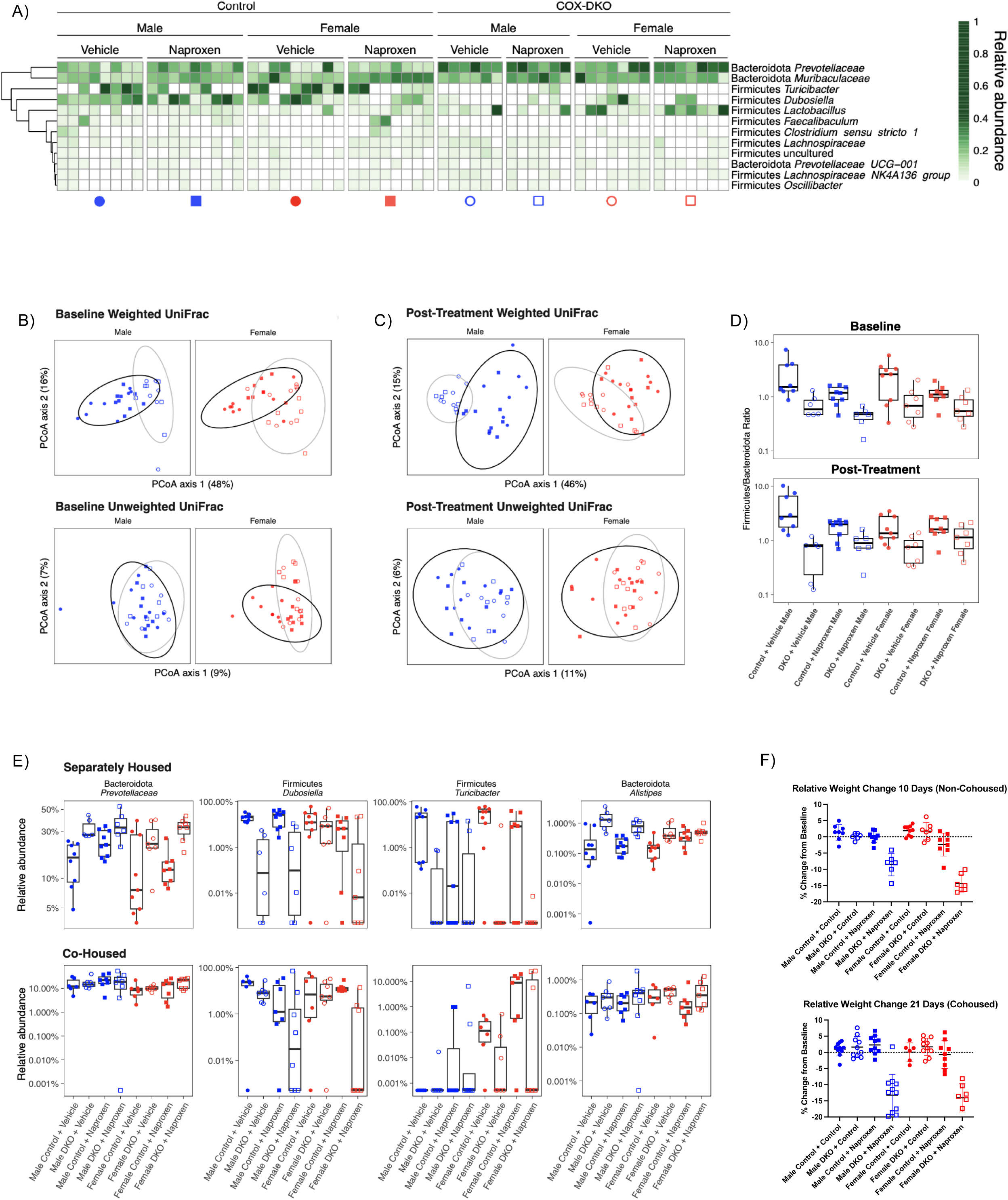
Divergent gut microbiome composition at baseline predisposes *Cox*-DKO mice to more severe enteropathy when treated with naproxen. *Cox*-DKO and *Cre^-/-^* control mice were treated with either control diet or naproxen diet (230 mg/kg) and allowed to feed *ad libitum* for either 10 days (separately housed by genotype) or 21 days (co-housed) prior to tissue collection; difference in duration was due to increased rate of naproxen-induced weight loss when mice were separately housed by genotype. n = 7-10 mice per group for entire figure. (**A**) Baseline bacterial taxa relative abundance heatmap including taxa at genus level or higher with an average relative abundance of at least 1% across all fecal samples. White tiles indicate no detection. (**B-C**) Beta diversity PCoA plots of bacterial UniFrac distances at baseline (**B**) and after 10 days of naproxen treatment (**C**). Ellipses denote 95% confidence intervals on genotypes: Control (black) and *Cox*-DKO (grey). (**D**) Boxplots of the ratio of Firmicutes to Bacteroidota relative abundances at baseline (top) and after 10 days of naproxen treatment (bottom). (**E**) Boxplots of differentially abundant taxa between genotypes at baseline. Top: separately housed animals. Bottom: co-housed animals. (**F**) Change in body weight relative to baseline in separately housed (10 days) vs co-housed (21 days) experiments.

The relative increase in Gram-negative relative to Gram-positive taxa in the *Cox*-DKO mice is consistent with previously published data on NSAID-induced enteropathy.^[51, 53, 54, 64–66]^ However, here we associate this result with prostaglandin suppression in the absence of enteropathy and, unlike prior acute studies with other NSAIDs, we do not observe dysbiosis with chronic dosing of naproxen alone. High-abundance genera or families that were distinct between *Cox*-DKOs and control mice at baseline include *Turicibacter* (males q=0.17, females q<0.05), *Dubosiella* (males q<0.05), *Prevotellaceae* (males q=0.06, females q<0.05), and *Alistipes* (males q<0.05, females q=0.1); however, these differential patterns are markedly dampened when littermate animals were not separated by genotype (all q>0.05), shifting the *Cox*-DKO gut microbiome composition towards that of the controls (Figure 4E). No taxa met the false discovery rate cutoff when littermates were not separated by genotype (Supplemental Figures 5A and 5B). This cohousing of the *Cox*-DKOs with controls also delayed the onset of naproxen-induced weight loss (Figure 4F), suggesting a protective role of commensal microbiota^[64, 66]^ from NSAID-associated enteropathy.

**Figure 5:**
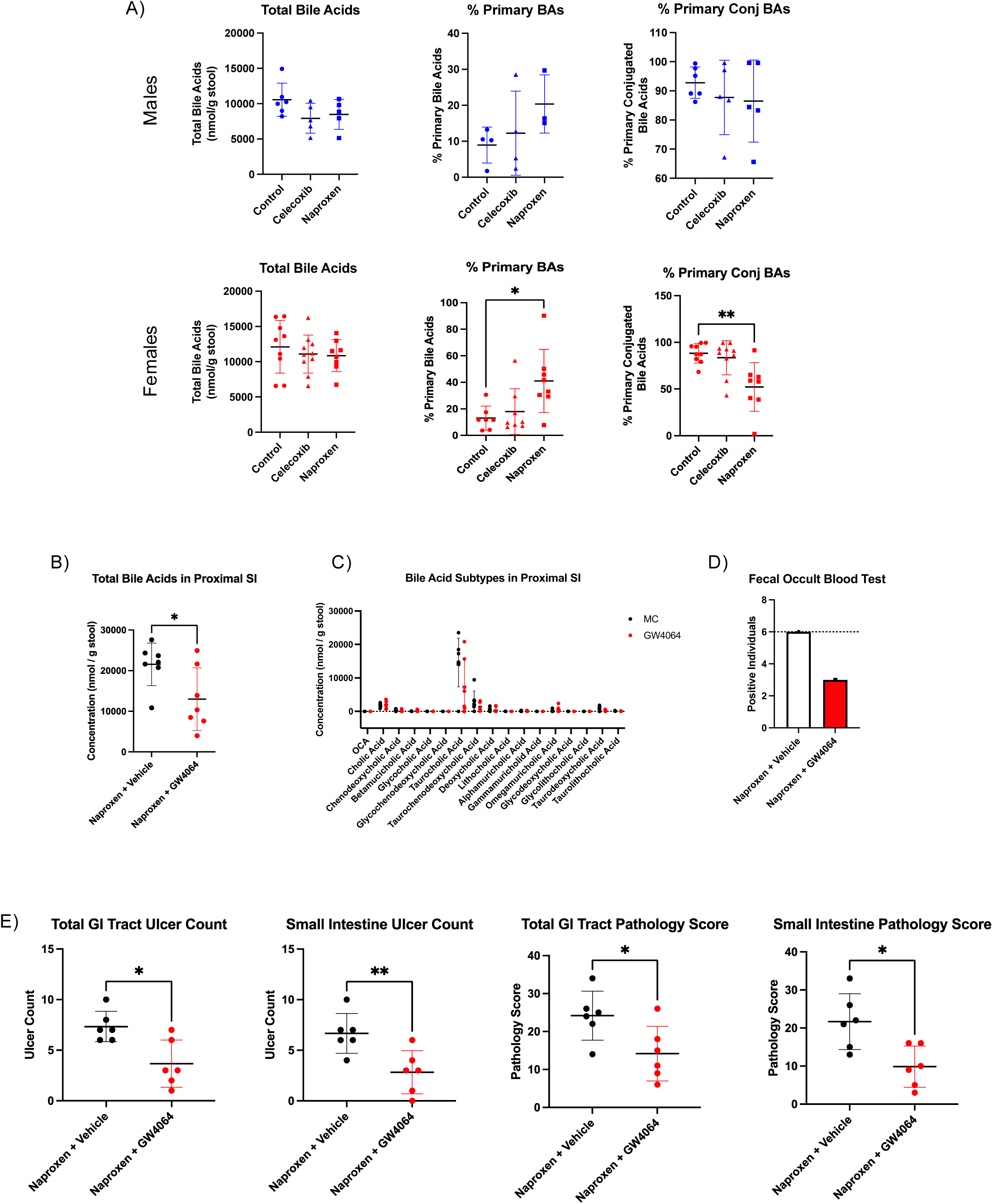
Distinct gut microbe-dependent bile acid conversions are observed with chronic naproxen treatment, and suppression of bile acid synthesis protects against naproxen-induced enteropathy in a COX-independent manner. (**A**) Wildtype C57BL/6J mice were treated with either control diet, celecoxib diet (100 mg/kg), or naproxen diet (230 mg/kg) and allowed to feed *ad libitum* for 3 weeks prior to tissue collection. n = 5-9 mice per group. Bile acids were measured in intestinal luminal content samples via UPLC. *p<0.05, **p<0.01 by unpaired t test. (**B-E**) *Cox*-DKO mice were fed naproxen diet (230 mg/kg) *ad libitum*, paired with either 30 mg/kg GW4064 or 0.5% methylcellulose (vehicle) by twice daily oral gavage for 10 days. n = 6 mice per group, females only. (**B**) Total bile acids in proximal small intestine luminal contents measured by UPLC. *p<0.05 by unpaired t test. (**C**) Individual bile acids in proximal small intestine luminal contents measured by UPLC. (**D**) Hemoccult test reported as number of mice that tested positive in each treatment group within the 10-day period. (**E**) Ulcer count for total GI tract vs small intestine only, as well as pathology score for total GI tract vs small intestine only. *p<0.05, **p<0.01 by unpaired t test.

We sought to detect bacteria-derived metabolites of functional relevance that might result from the dysbiosis consequent to *Cox* depletion. A targeted measurement of entero-protective short-chain fatty acids (Supplemental Figure 6A) and an untargeted screen of metabolites (Supplemental Figure 6B) failed to identify candidates. Similarly, *Cox* depletion did not result in an impact on immune cells, measured by flow cytometry, that could predispose the mice to exacerbate NSAID-induced enteropathy; proinflammatory patterns in myeloid cells that did not discriminate by genotype were only detected on naproxen challenge (Supplemental Figure 7). We also used an acute dose of indomethacin, an NSAID known to undergo enterohepatic recirculation, to address the possibility that the DKO-induced dysbiosis might influence bacterial glucuronidase activity to modify the duration of drug exposure^[55, 60, 67]^ (Supplemental Figure 8A). However, there was no impact of genotype (Supplemental Figure 8B).

Naproxen also undergoes enterohepatic recirculation, and intestinal bacteria may influence bile acid conjugation; deconjugation of primary bile acids has been shown to increase their toxicity to intestinal mucosa.^[66, 68]^ We observed that naproxen exposure in wildtype C57BL/6 mice resulted in decreased bile acid conjugation (Figure 5A). Bile acids activate the farnesoid X receptor (FXR), which down regulates CYP7A1 and thereby the hepatic conversion of cholesterol to primary bile acids. Here we showed that the FXR agonist, GW4064 suppressed total bile acid levels in the proximal small intestinal lumen relative to vehicle (0.5% methylcellulose) on a *Cox*-DKO background (Figure 5B). This suppression was largely driven by the most abundant bile acid subtype, taurocholic acid (Figure 5C). We then demonstrated that coadministration of the FXR agonist with naproxen protected against naproxen-induced GI bleeding (Figure 5D) and reduced ulcer count and pathology score (Figure 5E), and this protective effect was predominantly seen in the small intestine.

To address the possibility that the gut microbiome might influence NSAID-induced enteropathy in humans, we assessed the impact of coincident antibiotic treatment with NSAIDs on enteropathy in two human retrospective cohorts – the UK Biobank^[69]^ and All of Us Research Cohort Program.^[70]^ In both data sets, the incidence of NSAID-associated GI bleeding was greater when they were co-administered with antibiotics, potentially implicating the ablation of protective commensal bacteria.^[64, 66]^ (Figures 6A and 6B). For the UK Biobank population, the relative incidence of GI bleeding for patients taking NSAIDs alone vs in combination with antibiotics was 6.87% vs 9.48% (Figure 6A, p=1.9e^-35^). For the All of Us population, the relative incidence of GI bleeding for patients taking NSAIDs alone vs in combination with antibiotics was 4.10% vs 8.47% (Figure 6B, p<1.0e^-100^).

**Figure 6:**
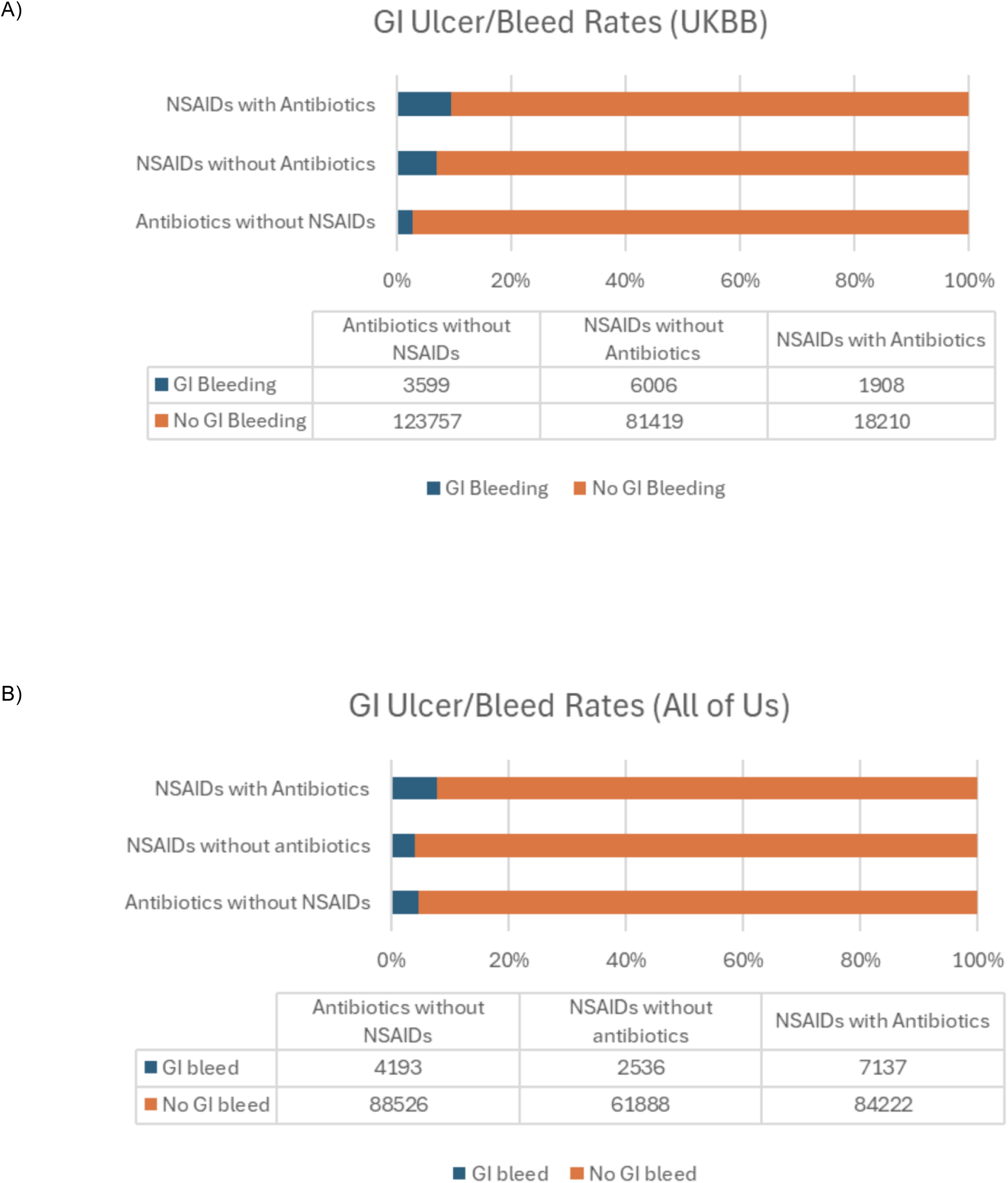
Coincident administration of NSAIDs and antibiotics in humans increases the risk of GI bleeding. Queries of GI ulceration and bleeding rates in human patients taking NSAIDs alone, antibiotics alone, or concurrent NSAIDs and antibiotics using UK Biobank (**A**) and All of Us (**B**). For the UK Bio Bank population, the relative incidence of GI bleeding for patients taking NSAIDs alone vs in combination with antibiotics was 6.87% vs 9.48% (p=1.9e^-35^); for the All of Us population, the relative incidence of GI bleeding for patients taking NSAIDs alone vs in combination with antibiotics was 4.10% vs 8.47% (p<1.0e^-100^); p-values calculated by Fisher’s exact test.

## Discussion

Although multiple mechanisms have been implicated in NSAID induced enteropathy, here we make the surprising observation that suppression of PG formation alone is of minimal, if any importance. Prolonged follow up of mice depleted postnatally of both COXs failed to result in enteropathy despite prostaglandin suppression. Furthermore, both the COX inhibitor, naproxen and its structural analog, phenylpropionic acid, devoid of COX inhibitory activity evoke enteropathy. Thus, the enteropathy evoked by clinical levels of naproxen exposure is largely independent of its ability to suppress PG production.

A second surprising observation was that although adding naproxen to COX depleted mice does not further suppress prostaglandin production, the severity of the NSAID induced enteropathy is markedly augmented when compared to its evocation in wildtype mice. Depletion or inhibition of the COX enzymes caused dysbiosis of the gut microbiome and, consistent with prior observations we observed a relative increase in Gram-negative taxa in the DKOs compared to control mice. While such a shift in the composition of the gut microbiota might have multiple effects of relevance to the augmentation of naproxen evoked enteropathy, we saw no impact on absorbed bacterial metabolites, host immune function, or drug metabolism.

Consistent with a previous report,^[66]^ we found that naproxen may have increased bile acid cytotoxicity by promoting the deconjugation of primary bile acids in wildtype mice. Thus, co-administration of an FXR agonist to the DKO mice significantly reduced the severity of naproxen evoked enteropathy, as previously reported in wildtype mice.^[71]^ Bile duct ligation also successfully prevents the formation of indomethacin-induced enteric ulcers in mice,^[51]^ further implicating bile acids as an important contributor to NSAID-induced enteropathy.

To seek clinical evidence connecting the dysbiosis caused by PG suppression with NSAID evoked enteropathy, we compared its incidence in patients with and without concomitant treatment with antibiotics. The higher rates of enteropathy in the UK Biobank and All of Us on the drug combination are consistent with the hypothesis that microbial ablation by the antibiotics removed commensals that limit NSAID evoked tissue injury, whether by topical irritation, bile acids, or other mechanisms. Postoperatively, patients often receive both treatments.

Although naproxen induced enteropathy appears more common and severe in females in the mice, this observation requires further investigation, given the sample sizes and very limited support in existing literature.^[52, 72, 73]^ Similarly, clinical trials of NSAIDs are heavily biased towards inclusion of women,^[74–76]^ and interrogation of a sex dependent incidence of NSAID induced enteropathy in humans has been similarly underpowered.^[77–80]^

In summary, PG suppression plays a trivial role in NSAID-induced enteropathy. However, *Cox* deletion causes dysbiosis of the gut microbiome that amplifies the enteropathic response to NSAIDs.

## Methods

### Sex as a Biological Variable

Both male and female mice and humans were included throughout all experiments and analyzed separately vs combined to account for sex as a biological variable.

### Animals

All mice were bred and maintained in our animal facility and fed *ad libitum* with standard chow diet (5010, Laboratory Autoclavable Rodent Diet, St. Louis, MO) unless placed on a special formulated diet for a particular study. Mice were kept under a 12-hour light / 12-hour dark cycle, with lights on at 7:00 AM and lights off at 7:00 PM. Male and female mice aged 10-12 weeks were used for all experiments, and data was always analyzed separately for each sex. Experimental protocols were reviewed and approved by the Institute for Animal Care and Use Committee at the University of Pennsylvania (Protocol #804445).

### Genotypes

All mice were on a C57BL/6J background. *Cox-1^fx/fx^* mice^[81]^ were gifted to us from the Herschman Lab at the University of California Los Angeles, and they were selectively bred with our tamoxifen-inducible *Cox-2^fx/fx^ CMV-Cre^+/-^* mice^[82]^ that we maintained in-house to yield the *Cox-1^fx/fx^ Cox-2^fx/fx^ CMV-Cre^+/-^* and *Cox-1^fx/fx^ Cox-2^fx/fx^ CMV-Cre^-/-^* mice that were used throughout these experiments for universal *Cox-1* and *-2* double-knockout (“*Cox*-DKO”) and *Cre^-/-^* controls derived from the same litter. Primer sequences for genotyping were:

*Cox-1* Flox Forward 1: 5’-TACCGCTGTCTCAGATTTTCCAGC-3’

*Cox-1* Flox Reverse 1: 5’-GTGGAGCTGAAGCTAGGAAACAGC-3’

*Cox-1* Flox Reverse 2: 5’-CCAGGCTTTACACTTTATGCTTCCG-3’

*Cox-2* Flox Forward: 5’-TGAGGCAGAAAGAGGTCCAGCCTT-3’

*Cox-2* Flox Reverse: 5’-ACCAATACTAGCTCAATAAGTGAC-3’

*CMV-Cre* Forward: 5’-CGATGCAACGAGTGATGAGG-3’

*CMV-Cre* Reverse: 5’-GCATTGCTGTCACTTGGTCGT-3’

### Chemicals

All chemicals and materials used were purchased from Sigma-Aldrich (St. Louis, MO) unless otherwise stated.

### *In Vivo* Study Designs

#### Study 1 Design: Longitudinal Tracking of Cox-DKO Mice

Baseline weight, feces, and urine were collected (prior to Cre-recombinase activation by tamoxifen.) Then mice of both *Cox*-DKO and Cre-negative genotypes were administered tamoxifen (200 mg/kg body weight per day for 5 consecutive days, dissolved in 85% corn oil / 15% ethanol solution) via oral gavage at approximately 10-12 weeks of age to induce genetic deletion of *Cox-1* and *Cox-2*. One month was allowed for tamoxifen washout, then weight and feces were collected weekly, and urine was collected monthly for 10 months (until the mice reached approximately one year of age.) Gastrointestinal tissues were harvested and prepared for histology. n = 14 *Cox*-DKO mice and 6 *Cre^-/-^* controls.

#### Study 2 Design: LPS Challenge on Cox-DKO Background

Mice of both genotypes were administered tamoxifen (200 mg/kg body weight per day for 5 consecutive days, dissolved in 85% corn oil / 15% ethanol solution) via oral gavage at approximately 10-12 weeks of age to induce genetic deletion of *Cox-1* and *Cox-2*. One month was allowed for tamoxifen washout. Then after baseline urine collection, mice received a single dose of LPS (1 mg/kg body weight dissolved in 1X PBS) by IP injection, and urine was collected overnight in metabolic cages. n = 7-9 mice per group.

#### Study 3 Design: Chronic Naproxen Exposure on Wildtype Background

Wildtype C57BL/6J mice were fed control diet (Purina Lab Chow, Envigo, Indianapolis, IN) *ad libitum* for 2 weeks, at which point baseline weight, feces, and urine were collected. Then cages were assigned either control diet or an identical diet with 1323 ppm naproxen or 550 ppm celecoxib dry-mixed into the formulation (Naproxen Custom Diet, Envigo, Indianapolis, IN) for 3 weeks. Weight and feces were collected each week, and urine was collected at the end of the 3-week period prior to tissue harvest. Gastrointestinal tissues were fixed for histology, and blood was collected for complete blood counts and serum protein measurements. n = 6-9 mice per group.

#### Study 4 Design: Chronic Naproxen Exposure on Cox-DKO Background

Mice of both genotypes were administered tamoxifen (200 mg/kg body weight per day for 5 consecutive days, dissolved in 85% corn oil / 15% ethanol solution) via oral gavage at approximately 10-12 weeks of age to induce genetic deletion of *Cox-1* and *Cox-2*. One month was allowed for tamoxifen washout, during which mice were fed control diet (Purina Lab Chow, Envigo, Indianapolis, IN) *ad libitum*. Then after baseline weight, feces, and urine collection, cages were assigned either control diet or an identical diet with 1323 ppm naproxen dry-mixed into the formulation (Naproxen Custom Diet, Envigo, Indianapolis, IN) for 3 weeks. Weight and feces were collected each week, and urine was collected at the end of the 3-week period before tissues were harvested. Gastrointestinal tissues were fixed for histology, and blood was collected for complete blood counts. This experiment was performed two slightly different ways to assess relative contribution of the gut microbiome and host immune system: 1) littermate mice of both genotypes cohoused together in the same cage for the entire duration of the experiment, and 2) littermate mice separated by genotype prior to tamoxifen administration; this version was terminated at 10 days rather than 21 days due to increased rate of weight loss in *Cox*-DKO + naproxen cages. n = 7-10 mice per group.

#### Study 5 Design: Chronic Phenylpropionic Acid Exposure on Cox-DKO Background

Mice of both genotypes were administered tamoxifen (200 mg/kg body weight per day for 5 consecutive days, dissolved in 85% corn oil / 15% ethanol solution) via oral gavage at approximately 10-12 weeks of age to induce genetic deletion of *Cox-1* and *Cox-2*. One month was allowed for tamoxifen washout. Then after baseline weight, feces, and urine collection, mice were fed a diet with 1323 ppm phenylpropionic acid dry-mixed into the formulation (PPA Custom Diet, Envigo, Indianapolis, IN) for 3 weeks. Weight and feces were collected each week, and urine was collected at the end of the 3-week period before tissues were harvested and prepared for histology. n = 6-9 mice per group.

#### Study 6 Design: FXR Agonist and Naproxen Coadministration on Cox-DKO Background

*Cox-1^fx/fx^ Cox-2^fx/fx^ CMV-Cre^+/-^* mice were administered tamoxifen (200 mg/kg body weight per day for 5 consecutive days, dissolved in 85% corn oil / 15% ethanol solution) via oral gavage at approximately 10-12 weeks of age to induce genetic deletion of *Cox-1* and *Cox-2*. One month was allowed for tamoxifen washout. All mice were fed 1323 ppm naproxen diet (Envigo, Indianapolis, IN) *ad libitum*, paired with either 30 mg/kg bodyweight GW4064 (BePharm Scientific, Arlington Heights, IL) or 0.5% methylcellulose (vehicle) by twice daily oral gavage. Feces, small intestine luminal contents, and gastrointestinal tissue were collected after 10 days of treatment. n = 6 mice per group, females only.

#### Study 7 Design: Microbiome-Dependent Elimination of Indomethacin

Wildtype C57BL/6J mice were administered either an antibiotic cocktail (1 g/L ampicillin, 0.2 g/L vancomycin, 1 g/L neomycin, 1 g/L metronidazole, and 4 g/L aspartame) or vehicle (4 g/L aspartame) in their drinking water for one week prior to receiving a single dose of indomethacin (10 mg/kg bodyweight dissolved in PEG400) via oral gavage, and urine was collected for the following 4 hours to measure indomethacin-glucuronide / indomethacin ratio. n = 4 mice per group. Additionally, separately housed *Cox*-DKO mice and Cre-negative controls were treated with a single dose of indomethacin (10 mg/kg bodyweight dissolved in PEG400) via oral gavage, and urine was collected for the following 4 hours to measure urinary indomethacin-glucuronide / indomethacin ratio. n = 10-11 mice per group.

### Real-Time Polymerase Chain Reaction Analysis of Gene Expression

Total RNA was isolated from lung and small intestine tissue samples using the Qiagen RNeasy Kit (Qiagen, Germantown, MD). Reverse transcription was performed using the Applied Biosystems High-Capacity cDNA Reverse Transcription Kit (Applied Biosystems, Waltham, MA). Real-time polymerase chain reaction was performed using ABI TaqMan primers and reagents on an ABI Prism 7500 Thermocycler according to the manufacturer’s instructions. All mRNA measurements were normalized to *Hprt* mRNA levels via the 2^-ΔΔCT^ method. The following TaqMan primers were used:

*Ptgs1/Cox-1*: Mm00477214_m1 (Life Tech / Invitrogen / ABI, Carlsbad, CA)

*Ptgs2/Cox-2*: Mm00478374_m1 (Life Tech / Invitrogen / ABI, Carlsbad, CA)

*Hprt*: Mm01545399_m1 (Life Tech / Invitrogen / ABI, Carlsbad, CA)

### Mass Spectrometric Analysis of Prostanoids

Urinary prostaglandin metabolites were measured by liquid chromatography with tandem mass spectrometry (LC-MS/MS) as previously described.^[83]^ Briefly, urine samples were collected using metabolic cages over 4-hour periods. Labeled internal standards of known concentrations for metabolites of PGE_2_, PGD_2_, PGI_2_, and TxA_2_ were spiked into urine samples prior to solid phase extraction (SPE). Then SPE was performed using Strata-X 33μm polymeric reversed phase cartridges (Phenomenex, Torrence, CA). Spectra were generated using a Waters AQUITY UPLC system. Peak area ratios of target analytes to internal standards were calculated using TargetLynx 4.1 software (Waters Corp., Milford, MA). Urinary prostanoid metabolite measurements were normalized to urinary creatinine measurements.

### Histological Analysis of Gastrointestinal Injury

All histology sectioning, staining, and analysis was performed at the Penn Vet Comparative Pathology Core (RRID:SCR_022438). Briefly, during dissection the stomach was isolated, and intestinal tissues were separated into 4 segments: proximal small intestine (∼duodenum), middle small intestine (∼jejunum), distal small intestine (∼ileum), and colon. The stomach was first injected with fixative (Methacarn Solution: 60% methanol, 30% chloroform, 10% acetic acid) to preserve its shape, placed into a 50 mL conical tube filled with fixative, and later transferred to a cassette. Intestinal segments were Swiss Rolled, placed into separate 50 mL conical tubes filled with fixative, and later transferred to cassettes. Samples were then transported to the core facility to be dehydrated with ethanol, embedded in paraffin, cut into 5 μm sections, and then stained with hematoxylin and eosin (H&E) staining. Slides were then evaluated by a trained, fully blinded veterinary pathologist with no knowledge of respective treatment or genotype identities among the samples. The qualitative / semi-quantitative scoring system included assessment of ulcer distribution (none, focal, or multifocal), number of ulcers, ulcer severity (superficial, up to muscularis mucosa, or transmural), presence of peritonitis / serositis, and inflammation (none, minimal, mild, moderate, or severe).

### Fecal Occult Blood Test

Hemoccult Sensa Rapid Test Kit (Beckman Coulter, Brea, CA) was used to detect blood in freshly collected stool samples on a weekly basis. Briefly, if hemoglobin is present in a fecal sample, the resulting peroxidase activity catalyzes the oxidation of alpha-guaiaconic acid (in the test-strip) by hydrogen peroxide (in the developer solution) to form a conjugated blue quinone compound. If the test-strip turns blue, then the fecal sample is positive for occult blood. Results were reported as positive or negative (not quantifiable beyond binary designation.)

### Complete Blood Count Analysis

Whole blood (> 100 μL per sample) was collected in K2EDTA-coated tubes to prevent clotting and briefly stored at room temperature. Samples were then immediately transported to the CHOP Translational Core Laboratory within 3-4 hours of collection to perform complete blood count (CBC) analysis using a Sysmex XT-2000iV Automated Hematology Analyzer.

### Serum Protein Measurements

The serum proteins and liver enzymes were measured by the CHOP Translational Core using the platform Roche Cobas c311, a clinical automated chemistry analyzer.

### 16S rRNA Gene Sequencing

#### Library Preparation (16S V1-V2 High Biomass)

Bacterial DNA was isolated from fecal and small intestine luminal content samples via DNeasy PowerSoil Pro Kit (Qiagen, Germantown, MD). Isolated samples were then submitted to the Penn CHOP Microbiome Core for all subsequent steps. Barcoded PCR primers annealing to the V1-V2 region of the 16S rRNA gene were used for library generation. PCR reactions were carried out in duplicate using Q5 High-Fidelity DNA Polymerase (NEB, Ipswich, MA). Each PCR reaction contained 0.5 µM of each primer, 0.34 U Q5 Polymerase, 1X Buffer, 0.2 mM dNTPs, and 5 µL DNA in a total volume of 50 µL. Cycling conditions were as follows: 1 cycle of 98°C for 1 minute; 20 cycles of 98°C for 10 seconds, 56°C for 20 seconds, and 72°C for 20 seconds; 1 cycle of 72°C for 8 minutes. After amplification, duplicate PCR reactions were pooled and then purified using a 1:1 volume of SPRI beads. DNA in each sample was then quantified using PicoGreen and pooled in equal molar amounts. The resulting library was sequenced on the Illumina MiSeq using 2×250 bp chemistry. Extraction blanks and DNA-free water were subjected to the same amplification and purification procedure to allow for empirical assessment of environmental and reagent contamination. Positive controls, consisting of eight artificial 16S gene fragments synthesized in gene blocks and combined in known abundances, were also included.

#### Bioinformatics Processing

QIIME2 version 2023.2.0^[84]^ was employed to process sequencing reads with DADA2 version 1.26.0^[85]^ to denoise reads and identify amplicon sequence variants (ASVs). ASVs were assigned to taxonomies by sequence comparison to the SILVA 138 database^[86]^, using a Naïve Bayes classifier implemented in scikit-bio^[87]^. MAFFT^[88]^ was used to build a phylogenetic tree for calculating UniFrac distances^[89, 90]^.

#### Statistical Analyses

Statistical analyses of ASV tables and diversity metrics were performed with R libraries. UniFrac distances were visualized with Principal Coordinates Analysis plots, and differences between study groups were assessed using PERMANOVA^[91]^ with the adonisplus R package, which utilizes the vegan adonis2 software. PERMANOVAs were run with 999 permutations, and cage effect was accounted for by restricted shuffling of samples between cages. Differential abundance of any taxon with an average abundance of at least 0.1% across all fecal samples was assessed by generalized linear mixed effects models on log10-transformed relative abundances. Cage number was included as a random effect and genotype as a fixed effect. Multiple tests were adjusted for false discovery rate using the Benjamini-Hochberg method.

### Nuclear Magnetic Resonance Spectroscopy to Measure Microbiome-Associated Metabolites

All NMR experiments were performed using a Bruker Avance III HD NMR Spectrometer fitted with a 3mm TXI probe (Bruker Biospin, Billerica, MA). 50-100 mg per fecal sample were homogenized in water (10 µL per mg feces) and spun down at 10,000 x g for 10 minutes at 4°C; supernatant was transferred to a new tube, and spin step was repeated. Then supernatant was passed through a Durapore-PVDF 0.22 µm centrifugal filter (Merck Millipore Ltd., Cork, IRL) spun at 12,000 x g for 4 minutes at 4°C. Samples (180 µL) were dissolved in 20 mL of buffer containing D2O (Cortecnet Corp. New York, NY) and 0.26 mM internal standard (4,4-dimethyl-4-silapentane-1-sulfonic acid/DSS, Cambridge Isotope Laboratory, Andover, MA). Briefly, the first transient of a NOESY experiment was used for acquiring 1-dimensional NMR data with water signal saturation by continuous irradiation during relaxation delay (1 s) and mixing time (0.1 s). Each spectrum was acquired using 1024 scans, 76 K data points, and 14 ppm spectral width. The FIDs were zero-filled to 128 K; 0.1 Hz of linear broadening was applied followed by Fourier transformation. Metabolite levels in the spectra were quantified using a targeted profiling technique via Chenomx profiler V8.0 (Edmonton, AB, Canada). Multivariate data analysis was performed using Simca-P 17.0 (Sartorius Stedim, Aubagne, France). Principal Component Analysis was used to check the quality of data, followed by supervised Orthogonal Partial Least Square – Discriminant Analysis (OPLS-DA). OPLS-DA model was judged using Q^2^(cum) (cross validated R^2^ generated by 7-fold cross validation technique) and CV-ANOVA p values (< 0.05 denotes significant model).

### Tissue Preparation and Flow Cytometry

To generate single cell suspension of splenocytes, freshly harvested murine spleen was placed on a 70 µm strainer and gently macerated through the filter with a 3 mL syringe plunger. Strainer was flushed with a total of 6 mL RPMI 1640 + 10% FBS, then cells were spun down at 500 x g for 5 minutes at room temperature, and supernatant was then removed. Pellet was resuspended in 5 mL RBC lysis buffer and incubated for 10 minutes at room temperature. Reaction was stopped by adding 10 mL PBS, and cells were spun down again at 500 x g for 5 minutes at room temperature. The RBC lysis – PBS wash cycle was repeated one more time until the pellet was no longer red. Pellet was then resuspended in 1 mL FACS buffer (PBS + 0.1% BSA). Cells were separated at a dilution of ∼1X10^6^/sample. Blood was prepared in the same manner, sans maceration step.

In preparation for staining, cells were pre-incubated with TruStain FcX™ PLUS (anti-mouse CD16/32) Antibody (BioLegend) and Zombie UV Fixable Viability dye (BioLegend). Two separate staining panels were used. The first panel included antibodies for CD172a, CD317, Ly6C, SiglecF, CD11b, CD64, CD3, CD19, NK1.1, CD11c, Ly6G, XCR1, MHCII/IA-IE, CD45, F4/80, and B220. The second panel included antibodies for CD19, TCRγδ, CD3, CD44, NK1.1, CD127, CD62L, CD8b, CD45, CD4, CD8a, GATA3, Helios, T-bet, RORγt, and FoxP3. Samples were fixed with Foxp3/Transcription Factor Staining Buffer Set (eBioscience). For transcription factors staining, cells were permeabilized and stained in the Foxp3/Transcription Factor Staining Buffer Set at 4°C. Samples were recorded on a Cytek Aurora (Cytek Biosciences) and analyzed using FlowJo. Details for gating strategies and analyses have been described previously.^[92]^

### LC-MS/MS Analysis of Indomethacin and its Metabolite (Indomethacin Acyl-β-D-Glucuronide)

Urinary indomethacin and indomethacin acyl-β-D-glucuronide were measured as described previously.^[55]^ Briefly, 40 µL d4-Indomethacin (1 ng/µL) was spiked into the samples and brought to 1 mL by adding 920 µL water. Just before loading the sample for solid phase extraction (SPE), 20 µL formic acid was added to the mixture. SPE was performed using Strata-X 33μm polymeric reversed phase cartridges (Phenomenex, 8B-S100-TAK) by employing following steps: 1 mL methanol was added, followed by 0.25 mL water. The sample was loaded, washed with 1 mL water, and eluted with 1 mL methanol. The sample was dried and reconstituted in 100 µL of 10% acetonitrile. Separation of the compounds was carried out using a Waters ACQUITY UPLC system with an ultra-performance liquid chromatography (UPLC) column, 2.1 × 150 mm with 1.7 μm particles (Waters ACQUITY UPLC CSH C18) coupled with Waters TQS triple quadrupole instrument operated in electrospray negative ion mode. The precursor to product ion mass transitions used for indomethacin and indomethacin acyl-β-D-glucuronide were *m/z* 356.1/312.1 and *m/z* 532.1/193.1 using d4-indomethancin as an internal standard (*m/z* 360.1/316.1). Ten-point calibration samples (100 ng/µL, 50 ng/µL, 25 ng/µL, 12.5 ng/µL, 6.25 ng/µL, 3.125 ng/µL, 1.56 ng/µL, 0.78 ng/µL, 0.39 ng/µL, 0.195 ng/µL) were used for measuring indomethacin and its metabolite. The ratio of indomethacin acyl-β-D-glucuronide to indomethacin was determined using standard curves prepared in pooled mouse urine that was free of indomethacin and its metabolite.

### Bile Acid Measurements

Bile acid quantifications were performed by the Microbial Culture & Metabolomics Core of the Penn CHOP Microbiome Program and the Center for Molecular Studies in Digestive & Liver Diseases (NIH P30DK050306) as described previously.^[93, 94]^ Briefly, luminal contents were collected neat into pre-weighed tubes from the proximal half of the small intestine, and each individual sample weight was recorded (between 25-100 mg per sample). Samples were stored at -80°C. Samples and sample weights were then submitted to the Microbial Culture & Metabolomics Core, and 16 different bile acids were quantified in each sample using a Waters ACQUITY Ultra-Performance Liquid Chromatography (UPLC) System with a Cortecs UPLC C-18+ 1.6 μm 2.1 x 50 mm column and a QDa single quadrupole mass detector. Samples were suspended in methanol (5 μL/mg stool), vortexed for 1 minute, and centrifuged twice at 13,000g for 5 minutes. Intestinal flushes were vortexed for 1 minute, and centrifuged twice at 13,000g for 5 minutes. The supernatant was transferred to a new tube, sealed, and stored at 4°C until analysis. The flow rate was 0.8 mL/min, the injection volume was 4 uL, the column temperature was 30°C, the sample temperature was 4°C, and the run time was 4 min per sample. Eluent A was 0.1% formic acid in water; eluent B was 0.1% formic acid in acetonitrile; the weak needle wash was 0.1% formic acid in water; the strong needle wash was 0.1% formic acid in acetonitrile; and the seal wash was 10% acetonitrile in water. The gradient is: initial flow 70% eluent A; linear gradient to 100% eluent B over 2.5 minutes; hold at 100% eluent B for 0.6 minutes; and linear gradient to 70% eluent A over 0.9 minutes. The mass detection channels were: +357.35 for chenodeoxycholic acid and deoxycholic acid; +359.25 for lithocholic acid; -407.5 for cholic, alphamuricholic, betamuricholic, gammamuricholic, and omegamuricholic acids; -432.5 for glycolithocholic acid; -448.5 for glycochenodeoxycholic and glycodeoxycholic acids; -464.5 for glycocholic acid; -482.5 for taurolithocholic acid; -498.5 for taurochenodeoxycholic and taurodeoxycholic acids; and -514.4 for taurocholic acid. Samples were quantified against standard curves of at least five points run in triplicate (chemicals obtained from Santa Cruz Biotechnology, Dallas, TX and Steraloids Inc., Newport, RI). Standard curves were run at the beginning and end of each metabolomics run. Blanks and standards were run every eight samples. Measurements were normalized to luminal content sample weights.

### NSAIDs-Antibiotics Queries in Human Databases

To study rates of gastrointestinal bleeding following NSAID treatment and how those rates differ with concomitant antibiotic use in a real patient population, we created lists of NSAIDs and antibiotics which we queried in both UK Biobank^[69]^ and All of Us^[70]^. We defined patients taking concomitant NSAIDs and antibiotics as those taking antibiotics within 5 days of being prescribed an NSAID, filtering out prescriptions for topical antibiotics. We then compared rates of a composite gastrointestinal phenotype as defined by one or more instances of peptic ulcer, gastrojejunal ulcer, gastric ulcer, or duodenal ulcer in the medical records of those who did and did not receive antibiotics. We also created a list of antibiotics that specifically target Gram-positive and Gram-negative bacteria and compared rates of gastrointestinal bleeding between those classes of antibiotics as well. All p-values were calculated by Fisher’s exact test.

### Statistics

All statistical analyses were performed using GraphPad Prism 9, QIIME / RStudio (for 16S sequencing analyses), or SciPy / Jupyter Notebook (for analyses in All of Us and UK Biobank). One-way ANOVA, multiple unpaired t test, Mantel-Cox log-rank test, Fisher’s exact test, PERMANOVA, Orthogonal Partial Least Square – Discriminant Analysis (OPLS-DA), and methods of FDR correction were all conducted as described in each figure legend. Wherever applicable, all data were plotted as mean ± SD.

### Study Approval

Experimental protocols were reviewed and approved by the Institute for Animal Care and Use Committee at the University of Pennsylvania under Protocol Number 804445. The UK Biobank research has been conducted using the UK Biobank Resource under Application Number 32133.

## Data Availability

All underlying data can be accessed in the “Supporting Data Values” file.

## Author Contributions

KB wrote the manuscript, generated the *Cox*-DKO mouse line, designed / executed all *in-vivo* experiments involving the transgenic mice, and directed collaborations regarding histology, flow cytometry, 16S sequencing, bile acids, metabolomics, and the human data. ER generated the chronic NSAID dosing model, supervised the project, acquired funding, and edited the manuscript. SG, USD, and RJ conducted all LC-MS/MS analyses. CH conducted the flow cytometry experiments. CAA performed all gastrointestinal pathology scoring assessments. CT and NW conducted all 16S sequencing analyses. KK performed queries in human databases. AS conducted all NMR analyses. MDR, KB, AW, KC, FDB, and GDW oversaw the above experiments and provided reagents. GAF supervised the project, acquired funding, and edited the manuscript.

## Acknowledgments

We would like to acknowledge Drs. Ian Henrich, Ronan Lordan, and Elizabeth Hennessy for assistance with editing, in addition to Drs. Soon Yew Tang, Katherine Theken, Tilo Grosser, Christoph Thaiss, Michael Silverman, Chris Lengner, Elliot Friedman, Harvey Herschman, Jia Nong, and Enrico Radaelli for technical support. We would also like to acknowledge the Penn Vet Comparative Pathology Core (RRID:SCR_022438) and the Microbial Culture & Metabolomics Core of the Penn CHOP Microbiome Program and the Center for Molecular Studies in Digestive & Liver Diseases (NIH P30DK050306) for their assistance with specialized readouts. The UK Biobank research has been conducted using the UK Biobank Resource under Application Number 32133. This work uses data provided by patients and collected by the NHS as part of their care and support. We gratefully acknowledge *All of Us* participants for their contributions, without whom this research would not have been possible. We also thank the National Institutes of Health’s *<u>All of Us</u>* <u>Research Program</u> for making available the participant data examined in this study.

## Figure Legends

**Supplemental Figure 1:**
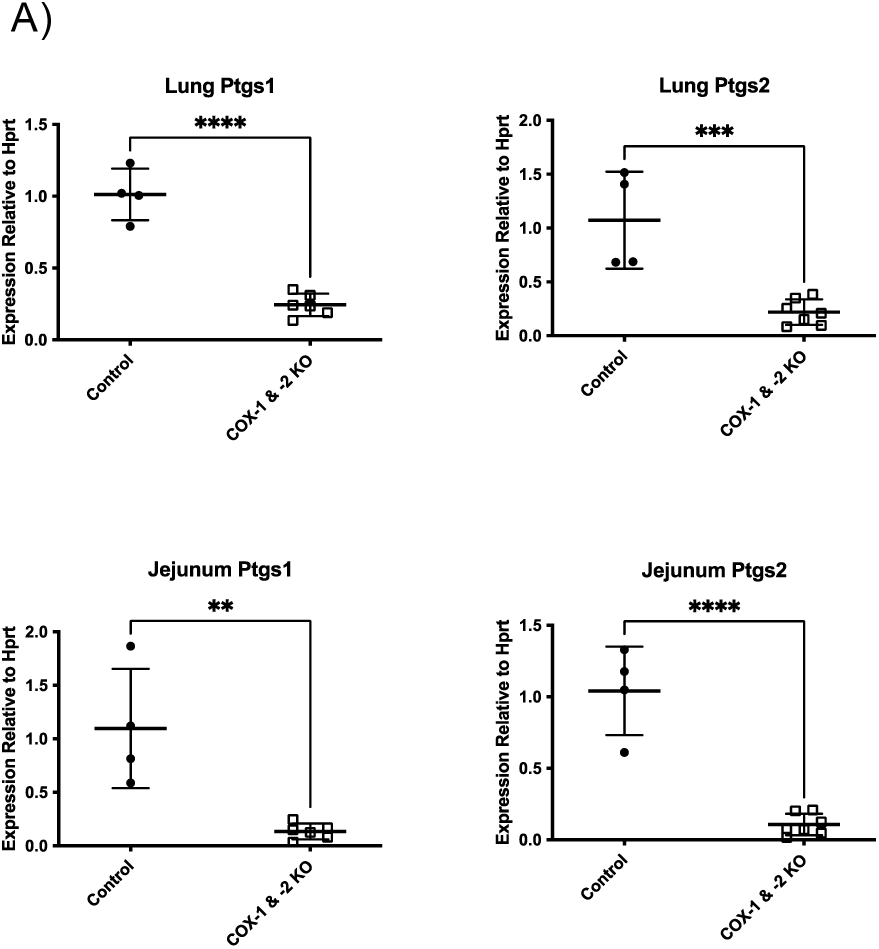
Confirmation of suppressed gene expression in lung and small intestine of inducible *Cox*-DKO mice. (**A**) Gene expression via qPCR for *Ptgs1* and *Ptgs2*, which encode COX-1 and COX-2, respectively, in both lung and small intestine tissue. **p<0.01, ***p<0.001, ****p<0.0001 by unpaired t test.

**Supplemental Figure 2:**
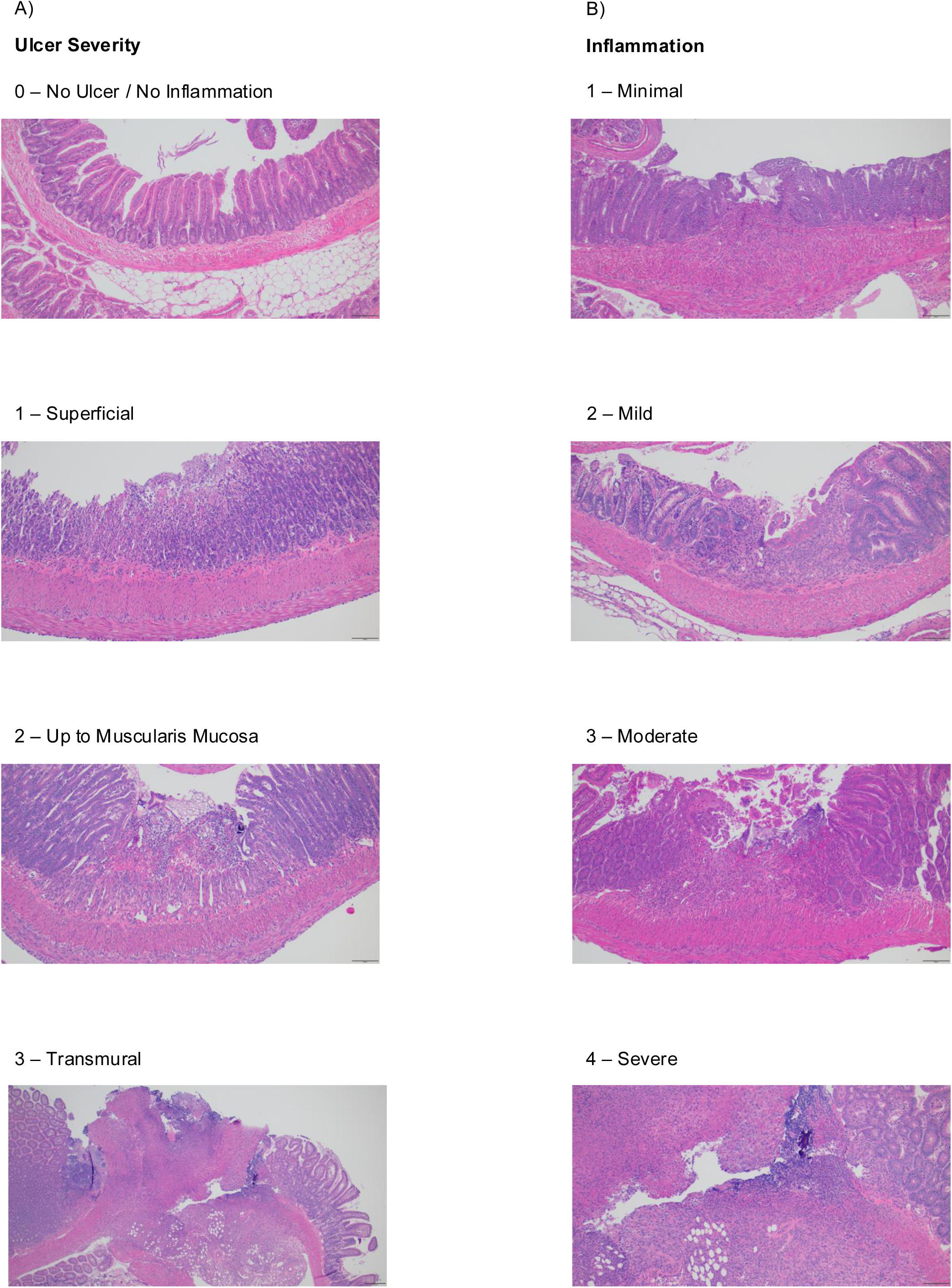
Representative small intestine histology images as scoring examples. H&E-stained murine small intestine samples collected after 3 weeks of naproxen diet. Representative pictures of ulcer severity (**A**) and inflammation (**B**) displayed increasing from top to bottom.

**Supplemental Figure 3:**
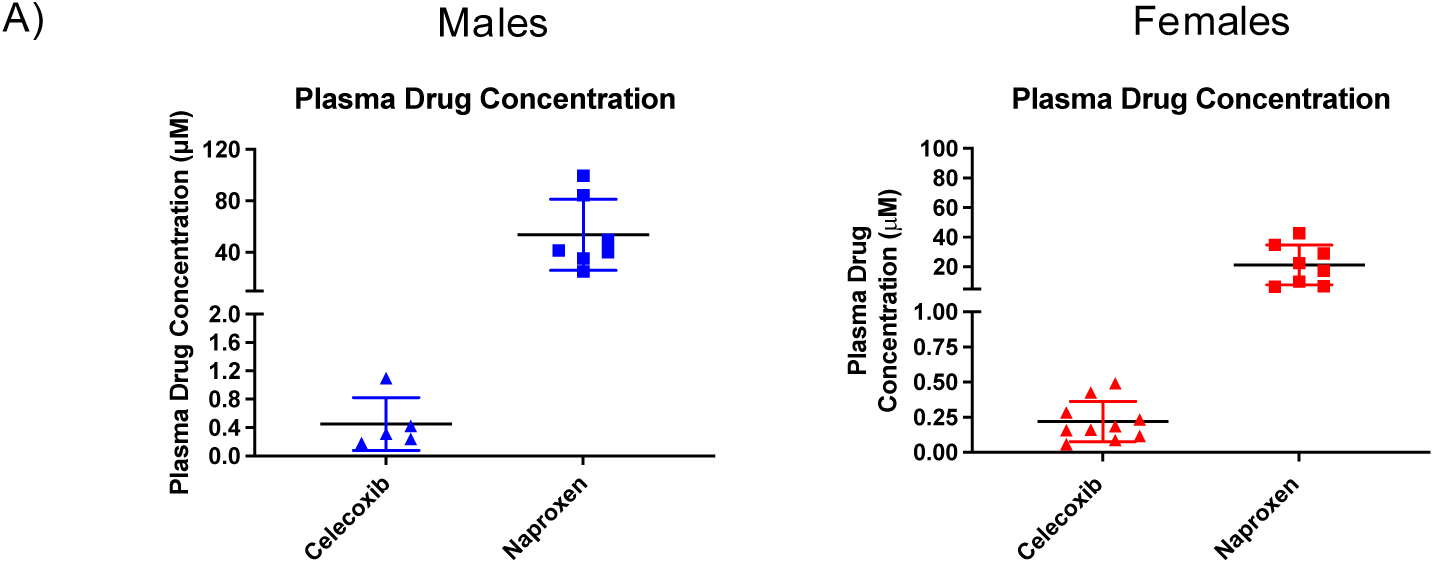
Confirmation of plasma drug concentrations in chronic NSAID dosing model. Wildtype C57BL/6J mice were treated with either control diet, celecoxib diet (100 mg/kg), or naproxen diet (230 mg/kg) and allowed to feed *ad libitum* for 3 weeks prior to tissue collection. (**A**) Plasma drug concentrations of celecoxib and naproxen measured by LC-MS/MS.

**Supplemental Figure 4:**
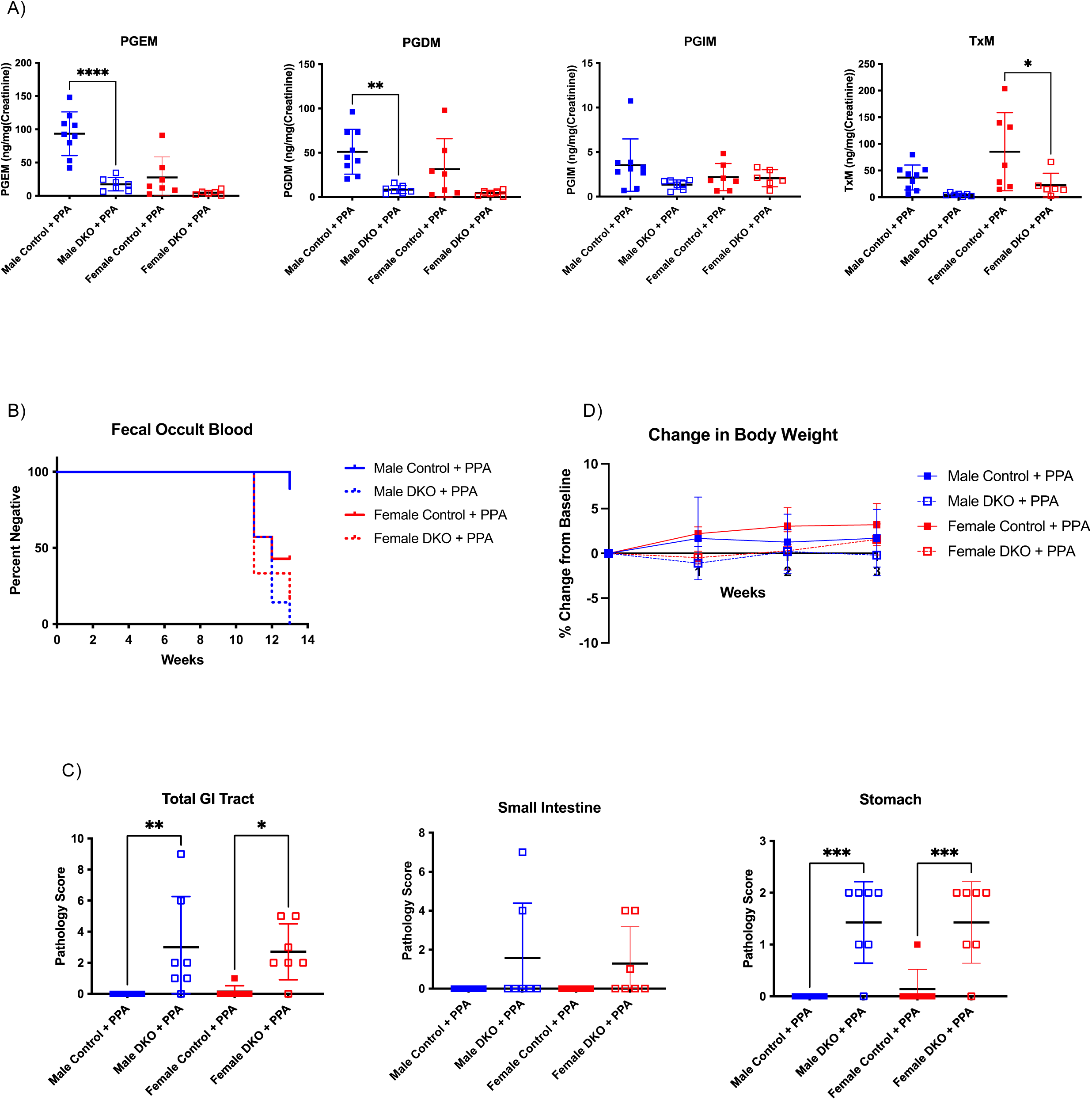
*Cox*-DKO mice develop gastrointestinal bleeding and lesions upon exposure to NSAID-analog phenylpropionic acid. *Cox*-DKO and *Cre^-/-^* control mice were treated with phenylpropionic acid diet (230 mg/kg) and allowed to feed *ad libitum* for 3 weeks prior to tissue collection. n = 6-9 mice per group for entire figure. (**A**) Urinary prostaglandin metabolites measured by LC-MS/MS. *p<0.05, **p<0.01, ****p<0.0001 by unpaired t test. (**B**) Weekly hemoccult test results plotted as a Kaplan-Meier curve for percentage of each group that tested negative for blood in the stool. Any individual that tested positive would be marked positive for that first week and all subsequent weeks, resembling a survival curve. (**C**) Pathology score for total GI tract, small intestine alone, and stomach alone. *p<0.05, **p<0.01, ***p<0.001 by unpaired t test. (**D**) Percent change in body weight relative to baseline body weight.

**Supplemental Figure 5:**
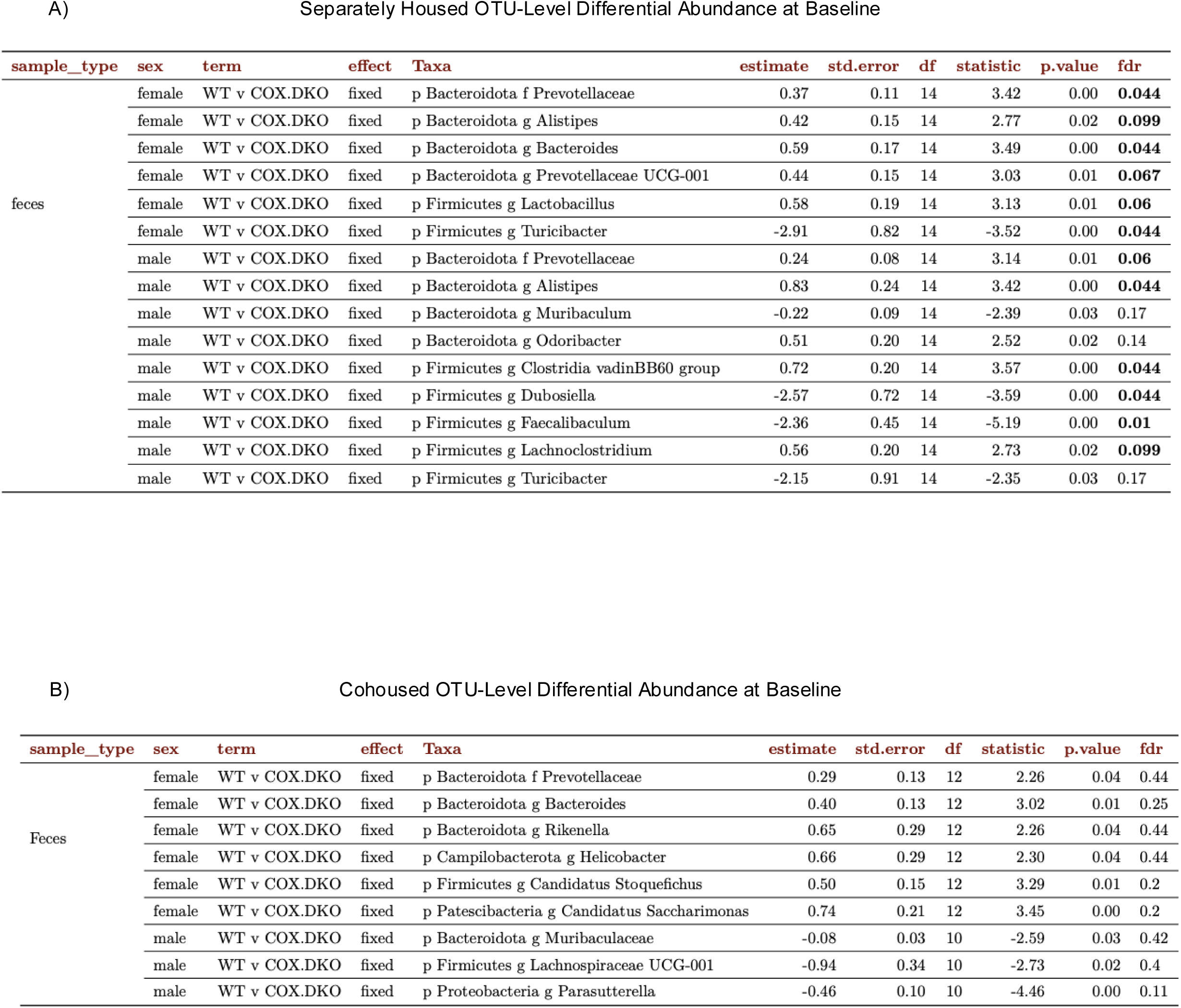
Distinct microbiome composition in *Cox*-DKO mice is masked by cohousing. OTU-level differential abundance at baseline for separately housed animals (**A**) and cohoused animals (**B**). Differential abundance of any taxon with an average abundance of at least 0.1% across all fecal samples was assessed by generalized linear mixed effects models on log10-transformed relative abundances. Multiple tests were adjusted for false discovery rate (FDR) using the Benjamini-Hochberg method. Any taxon with an FDR < 0.1 is displayed in bold text.

**Supplemental Figure 6:**
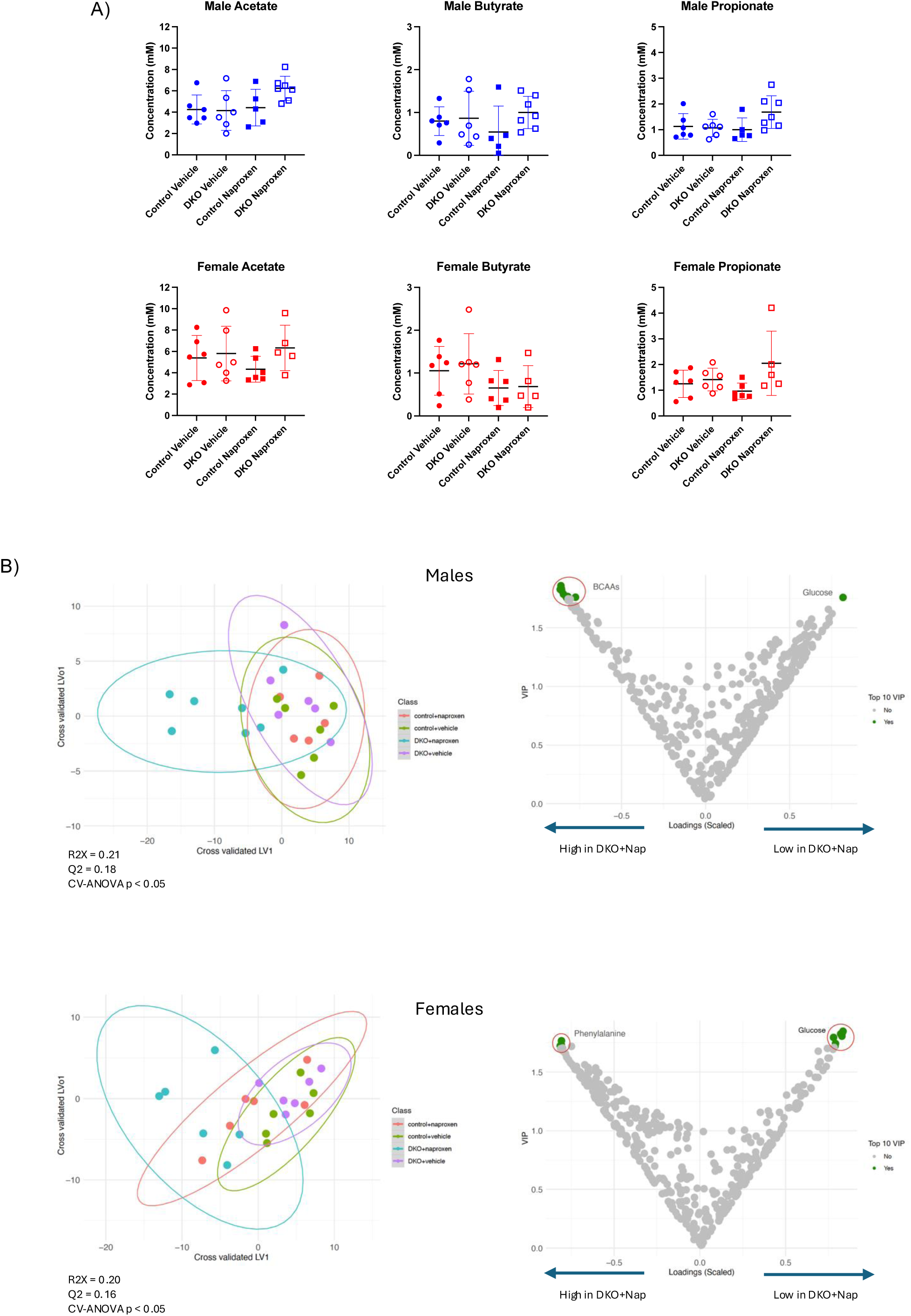
Targeted and untargeted metabolomics revealed no distinct fecal bacteria-derived metabolites of interest, apart from low glucose in the *Cox*-DKO mice treated with naproxen. Separately housed *Cox*-DKO and *Cre^-/-^* control mice were treated with either control diet or naproxen diet (230 mg/kg) and allowed to feed *ad libitum* for 10 days prior to tissue collection. n=5-7 mice per group for entire figure. (**A**) Measurements of fecal short-chain fatty acids (acetate, butyrate, and propionate) via targeted NMR. (**B**) Principal Component Analysis plots of fecal metabolites clustered by treatment group via untargeted NMR, paired with supervised Orthogonal Partial Least Square – Discriminant Analysis (OPLS-DA). *p<0.05 by CV-ANOVA.

**Supplemental Figure 7:**
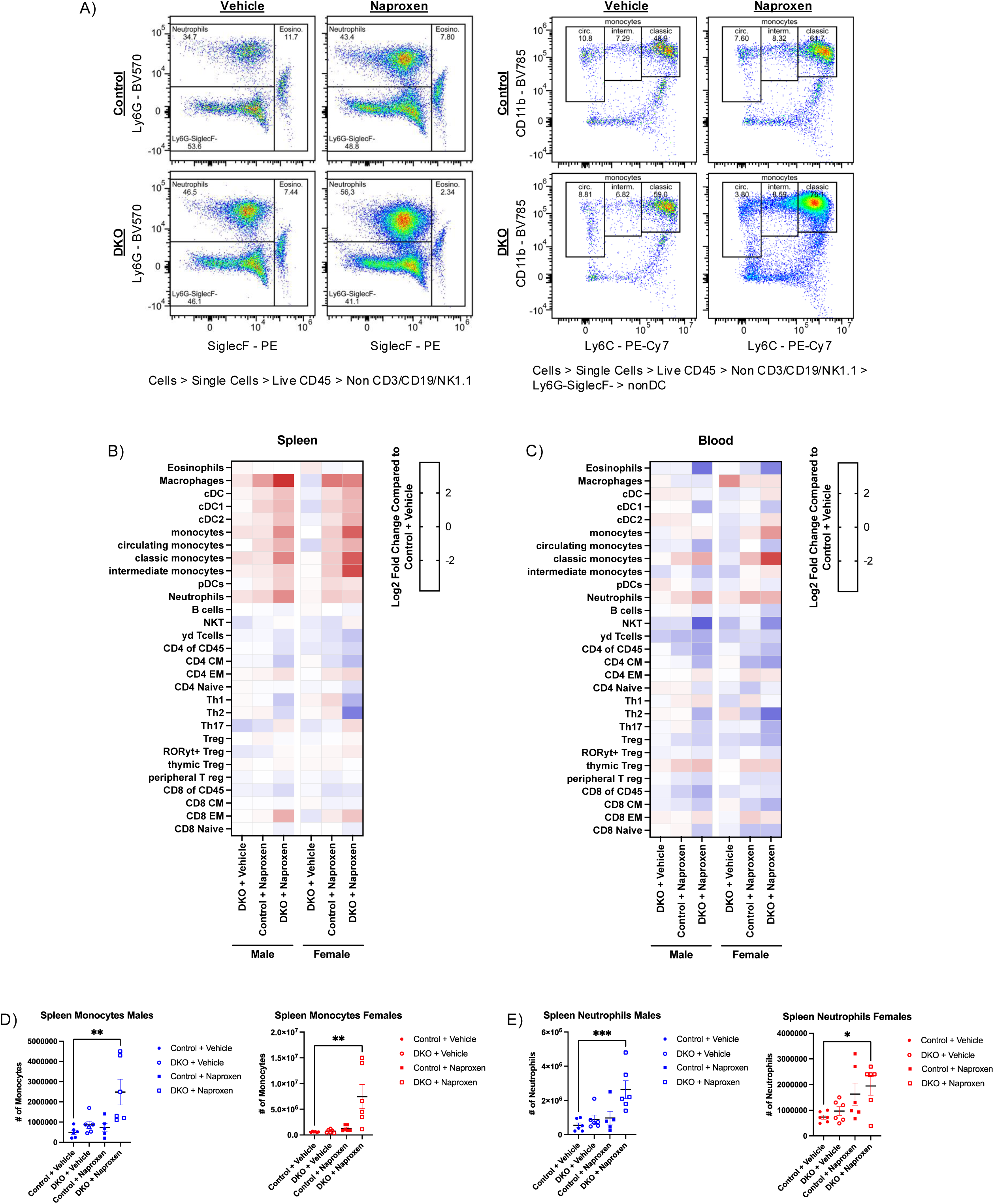
Divergent gut microbiome composition did not result in altered immune cell populations in *Cox*-DKOs at baseline. Separately housed *Cox*-DKO and *Cre^-/-^* control mice were treated with either control diet or naproxen diet (230 mg/kg) and allowed to feed *ad libitum* for 10 days prior to tissue collection. n=5-7 mice per group for entire figure. (**A**) Representative flow cytometry plots for monocytes and neutrophils. Heatmap summary of flow cytometry for spleen (**B**) and blood (**C**). Individual data points plotted for spleen monocytes (**D**) and spleen neutrophils (**E**). *p<0.05, **p<0.01, ***p<0.001 by one-way ANOVA.

**Supplemental Figure 8:**
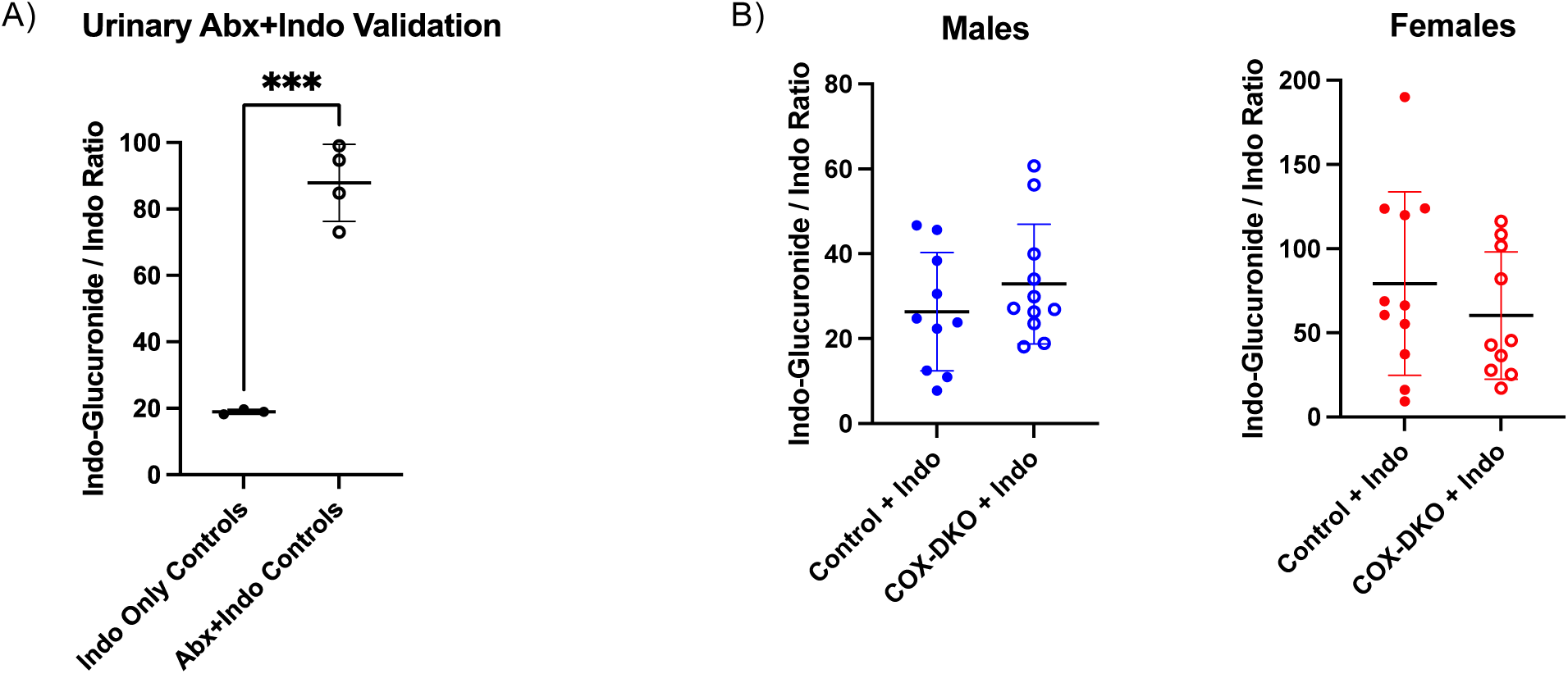
Baseline differences in microbiome composition between *Cox*-DKOs and controls did not result in differential drug elimination kinetics via deglucuronidation. (**A**) Wildtype C57BL/6J mice were administered either an antibiotic cocktail (1 g/L ampicillin, 0.2 g/L vancomycin, 1 g/L neomycin, 1 g/L metronidazole, and 4 g/L aspartame) or vehicle (4 g/L aspartame) in their drinking water for one week prior to receiving a single dose of indomethacin (10 mg/kg bodyweight dissolved in PEG400) via oral gavage, and urine was collected for the following 4 hours. Urinary indomethacin-glucuronide / indomethacin ratio measured by LC-MS/MS. n = 4 mice per group. ***p<0.001 by unpaired t test. (**B**) Separately housed *Cox*-DKO and *Cre^-/-^* control mice were treated with a single dose of indomethacin (10 mg/kg bodyweight dissolved in PEG400) via oral gavage, and urine was collected for the following 4 hours. Urinary indomethacin-glucuronide / indomethacin ratio measured by LC-MS/MS. n = 10-11 mice per group.

## References

1. Bhala, N., et al., Vascular and upper gastrointestinal effects of non-steroidal anti-inflammatory drugs: meta-analyses of individual participant data from randomised trials. Lancet, 2013. 382(9894): p. 769–79.

2. Paakkari, H., Epidemiological and financial aspects of the use of non-steroidal anti-inflammatory analgesics. Pharmacol Toxicol, 1994. 75 Suppl 2: p. 56–9.

3. Bjarnason, I., et al., Mechanisms of Damage to the Gastrointestinal Tract From Nonsteroidal Anti-Inflammatory Drugs. Gastroenterology, 2018. 154(3): p. 500–514.

4. Grosser, T., E. Ricciotti, and G.A. FitzGerald, The Cardiovascular Pharmacology of Nonsteroidal Anti-Inflammatory Drugs. Trends Pharmacol Sci, 2017. 38(8): p. 733–748.

5. Lanas, A. and F. Sopeña, Nonsteroidal anti-inflammatory drugs and lower gastrointestinal complications. Gastroenterol Clin North Am, 2009. 38(2): p. 333–52.

6. García-Rayado, G., M. Navarro, and A. Lanas, NSAID induced gastrointestinal damage and designing GI-sparing NSAIDs. Expert Rev Clin Pharmacol, 2018. 11(10): p. 1031–1043.

7. Martin, J.H., et al., Is cytochrome P450 2C9 genotype associated with NSAID gastric ulceration? Br J Clin Pharmacol, 2001. 51(6): p. 627–30.

8. Vonkeman, H.E., et al., Allele variants of the cytochrome P450 2C9 genotype in white subjects from The Netherlands with serious gastroduodenal ulcers attributable to the use of NSAIDs. Clin Ther, 2006. 28(10): p. 1670–6.

9. Ivey, K.J., Mechanisms of nonsteroidal anti-inflammatory drug-induced gastric damage. Actions of therapeutic agents. Am J Med, 1988. 84(2a): p. 41–8.

10. Overholt, B.F. and H.M. Pollard, Acetylsalicylic acid and ionic fluxes across the gastric mucosa of man. Gastroenterology, 1968. 54(4): p. 538–42.

11. Smith, B.M., et al., Permeability of the human gastric mucosa. Alteration by Acetylsalicylic acid and ethanol. N Engl J Med, 1971. 285(13): p. 716–21.

12. Ivey, K.J., S. Morrison, and C. Gray, Effect of salicylates on the gastric mucosal barrier in man. J Appl Physiol, 1972. 33(1): p. 81–5.

13. Rees, W.D., et al., The role of histamine receptors in the pathophysiology of gastric mucosal damage. Gastroenterology, 1977. 72(1): p. 67–71.

14. Kauffman, G.L., Jr. and M.I. Grossman, Prostaglandin and cimetidine inhibit the formation of ulcers produced by parenteral salicylates. Gastroenterology, 1978. 75(6): p. 1099–1102.

15. Daneshmend, T.K., et al., Abolition by omeprazole of aspirin induced gastric mucosal injury in man. Gut, 1990. 31(5): p. 514–7.

16. Yeomans, N.D., et al., A comparison of omeprazole with ranitidine for ulcers associated with nonsteroidal antiinflammatory drugs. Acid Suppression Trial: Ranitidine versus Omeprazole for NSAID-associated Ulcer Treatment (ASTRONAUT) Study Group. N Engl J Med, 1998. 338(11): p. 719–26.

17. Wallace, J.L., NSAID gastropathy and enteropathy: distinct pathogenesis likely necessitates distinct prevention strategies. Br J Pharmacol, 2012. 165(1): p. 67–74.

18. Vane, J.R., Inhibition of prostaglandin synthesis as a mechanism of action for aspirin-like drugs. Nat New Biol, 1971. 231(25): p. 232–5.

19. Ricciotti, E. and G.A. FitzGerald, Prostaglandins and inflammation. Arterioscler Thromb Vasc Biol, 2011. 31(5): p. 986–1000.

20. Smith, W.L. and D.L. Dewitt, Prostaglandin endoperoxide H synthases-1 and -2. Adv Immunol, 1996. 62: p. 167–215.

21. Jackson, B.A., et al., Effects of transforming growth factor beta and interleukin-1 beta on expression of cyclooxygenase 1 and 2 and phospholipase A2 mRNA in lung fibroblasts and endothelial cells in culture. Biochem Biophys Res Commun, 1993. 197(3): p. 1465–74.

22. Smith, C.J., et al., Differentiation of monocytoid THP-1 cells with phorbol ester induces expression of prostaglandin endoperoxide synthase-1 (COX-1). Biochem Biophys Res Commun, 1993. 192(2): p. 787–93.

23. Brannon, T.S., et al., Prostacyclin synthesis in ovine pulmonary artery is developmentally regulated by changes in cyclooxygenase-1 gene expression. J Clin Invest, 1994. 93(5): p. 2230–5.

24. Samet, J.M., et al., Selective induction of prostaglandin G/H synthase I by stem cell factor and dexamethasone in mast cells. J Biol Chem, 1995. 270(14): p. 8044–9.

25. Yamagata, K., et al., Expression of a mitogen-inducible cyclooxygenase in brain neurons: regulation by synaptic activity and glucocorticoids. Neuron, 1993. 11(2): p. 371–86.

26. Harris, R.C., et al., Cyclooxygenase-2 is associated with the macula densa of rat kidney and increases with salt restriction. J Clin Invest, 1994. 94(6): p. 2504–10.

27. Walenga, R.W., et al., Constitutive expression of prostaglandin endoperoxide G/H synthetase (PGHS)-2 but not PGHS-1 in hum an tracheal epithelial cells in vitro. Prostaglandins, 1996. 52(5): p. 341–59.

28. Zimmermann, K.C., et al., Constitutive cyclooxygenase-2 expression in healthy human and rabbit gastric mucosa. Mol Pharmacol, 1998. 54(3): p. 536–40.

29. MacNaughton, W.K. and K. Cushing, Role of constitutive cyclooxygenase-2 in prostaglandin-dependent secretion in mouse colon in vitro. J Pharmacol Exp Ther, 2000. 293(2): p. 539–44.

30. Whittle, B.J., Temporal relationship between cyclooxygenase inhibition, as measured by prostacyclin biosynthesis, and the gastrointestinal damage induced by indomethacin in the rat. Gastroenterology, 1981. 80(1): p. 94–8.

31. Moncada, S., et al., An enzyme isolated from arteries transforms prostaglandin endoperoxides to an unstable substance that inhibits platelet aggregation. Nature, 1976. 263(5579): p. 663–5.

32. Tanaka, A., et al., Role of cyclooxygenase (COX)-1 and COX-2 inhibition in nonsteroidal anti-inflammatory drug-induced intestinal damage in rats: relation to various pathogenic events. J Pharmacol Exp Ther, 2002. 303(3): p. 1248–54.

33. Sigthorsson, G., et al., COX-1 and 2, intestinal integrity, and pathogenesis of nonsteroidal anti-inflammatory drug enteropathy in mice. Gastroenterology, 2002. 122(7): p. 1913–23.

34. Tanaka, A., et al., Up-regulation of cyclooxygenase-2 by inhibition of cyclooxygenase-1: a key to nonsteroidal anti-inflammatory drug-induced intestinal damage. J Pharmacol Exp Ther, 2002. 300(3): p. 754–61.

35. Hotz-Behofsits, C., et al., Role of COX-2 in nonsteroidal anti-inflammatory drug enteropathy in rodents. Scand J Gastroenterol, 2010. 45(7-8): p. 822–7.

36. Wallace, J.L. and P.R. Devchand, Emerging roles for cyclooxygenase-2 in gastrointestinal mucosal defense. Br J Pharmacol, 2005. 145(3): p. 275–82.

37. Reuter, B.K., et al., Exacerbation of inflammation-associated colonic injury in rat through inhibition of cyclooxygenase-2. J Clin Invest, 1996. 98(9): p. 2076–85.

38. Mizuno, H., et al., Induction of cyclooxygenase 2 in gastric mucosal lesions and its inhibition by the specific antagonist delays healing in mice. Gastroenterology, 1997. 112(2): p. 387–97.

39. Masferrer, J.L., P.C. Isakson, and K. Seibert, Cyclooxygenase-2 inhibitors: a new class of anti-inflammatory agents that spare the gastrointestinal tract. Gastroenterol Clin North Am, 1996. 25(2): p. 363–72.

40. Morham, S.G., et al., Prostaglandin synthase 2 gene disruption causes severe renal pathology in the mouse. Cell, 1995. 83(3): p. 473–82.

41. Dinchuk, J.E., et al., Renal abnormalities and an altered inflammatory response in mice lacking cyclooxygenase II. Nature, 1995. 378(6555): p. 406–9.

42. Seibert, K., et al., Pharmacological and biochemical demonstration of the role of cyclooxygenase 2 in inflammation and pain. Proc Natl Acad Sci U S A, 1994. 91(25): p. 12013–7.

43. Bombardier, C., et al., Comparison of upper gastrointestinal toxicity of rofecoxib and naproxen in patients with rheumatoid arthritis. VIGOR Study Group. N Engl J Med, 2000. 343(21): p. 1520–8, 2 p following 1528.

44. Silverstein, F.E., et al., Gastrointestinal toxicity with celecoxib vs nonsteroidal anti-inflammatory drugs for osteoarthritis and rheumatoid arthritis: the CLASS study: A randomized controlled trial. Celecoxib Long-term Arthritis Safety Study. Jama, 2000. 284(10): p. 1247–55.

45. Iñiguez, M.A., et al., Detection of COX-1 and COX-2 isoforms in synovial fluid cells from inflammatory joint diseases. Br J Rheumatol, 1998. 37(7): p. 773–8.

46. Lichtenberger, L.M., et al., Non-steroidal anti-inflammatory drugs (NSAIDs) associate with zwitterionic phospholipids: insight into the mechanism and reversal of NSAID-induced gastrointestinal injury. Nat Med, 1995. 1(2): p. 154–8.

47. Lichtenberger, L.M., et al., NSAID injury to the gastrointestinal tract: evidence that NSAIDs interact with phospholipids to weaken the hydrophobic surface barrier and induce the formation of unstable pores in membranes. J Pharm Pharmacol, 2006. 58(11): p. 1421–8.

48. Manrique-Moreno, M., et al., The membrane-activity of Ibuprofen, Diclofenac, and Naproxen: a physico-chemical study with lecithin phospholipids. Biochim Biophys Acta, 2009. 1788(6): p. 1296–303.

49. Somasundaram, S., et al., Mitochondrial damage: a possible mechanism of the “topical” phase of NSAID induced injury to the rat intestine. Gut, 1997. 41(3): p. 344–53.

50. Somasundaram, S., et al., Uncoupling of intestinal mitochondrial oxidative phosphorylation and inhibition of cyclooxygenase are required for the development of NSAID-enteropathy in the rat. Aliment Pharmacol Ther, 2000. 14(5): p. 639–50.

51. Kent, T.H., R.M. Cardelli, and F.W. Stamler, Small intestinal ulcers and intestinal flora in rats given indomethacin. Am J Pathol, 1969. 54(2): p. 237–49.

52. Robert, A. and T. Asano, Resistance of germfree rats to indomethacin-induced intestinal lesions. Prostaglandins, 1977. 14(2): p. 333–41.

53. Hagiwara, M., et al., Role of unbalanced growth of gram-negative bacteria in ileal ulcer formation in rats treated with a nonsteroidal anti-inflammatory drug. J Med Invest, 2004. 51(1-2): p. 43–51.

54. Wallace, J.L., et al., Proton pump inhibitors exacerbate NSAID-induced small intestinal injury by inducing dysbiosis. Gastroenterology, 2011. 141(4): p. 1314-22, 1322.e1-5.

55. Liang, X., et al., Bidirectional interactions between indomethacin and the murine intestinal microbiota. Elife, 2015. 4: p. e08973.

56. Uejima, M., et al., Role of intestinal bacteria in ileal ulcer formation in rats treated with a nonsteroidal antiinflammatory drug. Microbiol Immunol, 1996. 40(8): p. 553–60.

57. Maseda, D. and E. Ricciotti, NSAID-Gut Microbiota Interactions. Front Pharmacol, 2020. 11: p. 1153.

58. Reuter, B.K., N.M. Davies, and J.L. Wallace, Nonsteroidal anti-inflammatory drug enteropathy in rats: role of permeability, bacteria, and enterohepatic circulation. Gastroenterology, 1997. 112(1): p. 109–17.

59. Zhou, Y., et al., Effect of indomethacin on bile acid-phospholipid interactions: implication for small intestinal injury induced by nonsteroidal anti-inflammatory drugs. Am J Physiol Gastrointest Liver Physiol, 2010. 298(5): p. G722–31.

60. Saitta, K.S., et al., Bacterial β-glucuronidase inhibition protects mice against enteropathy induced by indomethacin, ketoprofen or diclofenac: mode of action and pharmacokinetics. Xenobiotica, 2014. 44(1): p. 28–35.

61. Reese, J., et al., Coordinated regulation of fetal and maternal prostaglandins directs successful birth and postnatal adaptation in the mouse. Proc Natl Acad Sci U S A, 2000. 97(17): p. 9759–64.

62. Stanfield, K.M., et al., Expression of cyclooxygenase-2 in embryonic and fetal tissues during organogenesis and late pregnancy. Birth Defects Res A Clin Mol Teratol, 2003. 67(1): p. 54–8.

63. Fournel, S. and J. Caldwell, The metabolic chiral inversion of 2-phenylpropionic acid in rat, mouse and rabbit. Biochem Pharmacol, 1986. 35(23): p. 4153–9.

64. Kinouchi, T., et al., Culture supernatants of Lactobacillus acidophilus and Bifidobacterium adolescentis repress ileal ulcer formation in rats treated with a nonsteroidal antiinflammatory drug by suppressing unbalanced growth of aerobic bacteria and lipid peroxidation. Microbiol Immunol, 1998. 42(5): p. 347–55.

65. Koga, H., et al., Experimental enteropathy in athymic and euthymic rats: synergistic role of lipopolysaccharide and indomethacin. Am J Physiol, 1999. 276(3): p. G576–82.

66. Syer, S.D., et al., NSAID enteropathy and bacteria: a complicated relationship. J Gastroenterol, 2015. 50(4): p. 387–93.

67. LoGuidice, A., et al., Pharmacologic targeting of bacterial β-glucuronidase alleviates nonsteroidal anti-inflammatory drug-induced enteropathy in mice. J Pharmacol Exp Ther, 2012. 341(2): p. 447–54.

68. Hofmann, A.F., The continuing importance of bile acids in liver and intestinal disease. Arch Intern Med, 1999. 159(22): p. 2647–58.

69. Bycroft, C., et al., The UK Biobank resource with deep phenotyping and genomic data. Nature, 2018. 562(7726): p. 203–209.

70. Murray, J., The “All of Us” Research Program. N Engl J Med, 2019. 381(19): p. 1884.

71. Fiorucci, S., et al., Activation of the farnesoid-X receptor protects against gastrointestinal injury caused by non-steroidal anti-inflammatory drugs in mice. Br J Pharmacol, 2011. 164(8): p. 1929–38.

72. Chillingworth, N.L., S.G. Morham, and L.F. Donaldson, Sex differences in inflammation and inflammatory pain in cyclooxygenase-deficient mice. Am J Physiol Regul Integr Comp Physiol, 2006. 291(2): p. R327–34.

73. Craft, R.M., K.A. Hewitt, and S.C. Britch, Antinociception produced by nonsteroidal anti-inflammatory drugs in female vs male rats. Behav Pharmacol, 2021. 32(2&3): p. 153–169.

74. Solomon, D.H., et al., Differences in Safety of Nonsteroidal Antiinflammatory Drugs in Patients With Osteoarthritis and Patients With Rheumatoid Arthritis: A Randomized Clinical Trial. Arthritis Rheumatol, 2018. 70(4): p. 537–546.

75. Collantes, E., et al., A multinational randomized, controlled, clinical trial of etoricoxib in the treatment of rheumatoid arthritis [ISRCTN25142273]. BMC Fam Pract, 2002. 3: p. 10.

76. Nissen, S.E., et al., Cardiovascular Safety of Celecoxib, Naproxen, or Ibuprofen for Arthritis. N Engl J Med, 2016. 375(26): p. 2519–29.

77. Lee, S.P., et al., Effect of Nonsteroidal Anti-inflammatory Agents on Small Intestinal Injuries as Evaluated by Capsule Endoscopy. Dig Dis Sci, 2021. 66(8): p. 2724–2731.

78. Maiden, L., et al., A quantitative analysis of NSAID-induced small bowel pathology by capsule enteroscopy. Gastroenterology, 2005. 128(5): p. 1172–8.

79. Goldstein, J.L., et al., Video capsule endoscopy to prospectively assess small bowel injury with celecoxib, naproxen plus omeprazole, and placebo. Clin Gastroenterol Hepatol, 2005. 3(2): p. 133–41.

80. Edogawa, S., et al., Sex differences in NSAID-induced perturbation of human intestinal barrier function and microbiota. Faseb j, 2018. 32(12): p. fj201800560R.

81. Mitchell, J.A., et al., Cell-Specific Gene Deletion Reveals the Antithrombotic Function of COX1 and Explains the Vascular COX1/Prostacyclin Paradox. Circ Res, 2019. 125(9): p. 847–854.

82. Yu, Z., et al., Disruption of the 5-lipoxygenase pathway attenuates atherogenesis consequent to COX-2 deletion in mice. Proc Natl Acad Sci U S A, 2012. 109(17): p. 6727–32.

83. Song, W.L., et al., Noninvasive assessment of the role of cyclooxygenases in cardiovascular health: a detailed HPLC/MS/MS method. Methods Enzymol, 2007. 433: p. 51–72.

84. Bolyen, E., et al., Reproducible, interactive, scalable and extensible microbiome data science using QIIME 2. Nat Biotechnol, 2019. 37(8): p. 852–857.

85. Callahan, B.J., et al., DADA2: High-resolution sample inference from Illumina amplicon data. Nat Methods, 2016. 13(7): p. 581–3.

86. Quast, C., et al., The SILVA ribosomal RNA gene database project: improved data processing and web-based tools. Nucleic Acids Res, 2013. 41(Database issue): p. D590–6.

87. Bokulich, N.A., et al., Optimizing taxonomic classification of marker-gene amplicon sequences with QIIME 2’s q2-feature-classifier plugin. Microbiome, 2018. 6(1): p. 90.

88. Katoh, K. and D.M. Standley, MAFFT multiple sequence alignment software version 7: improvements in performance and usability. Mol Biol Evol, 2013. 30(4): p. 772–80.

89. Lozupone, C. and R. Knight, UniFrac: a new phylogenetic method for comparing microbial communities. Appl Environ Microbiol, 2005. 71(12): p. 8228–35.

90. Lozupone, C.A., et al., Quantitative and qualitative beta diversity measures lead to different insights into factors that structure microbial communities. Appl Environ Microbiol, 2007. 73(5): p. 1576–85.

91. Anderson, M.J., A new method for non-parametric multivariate analysis of variance. Austral Ecology, 2001. 26(1): p. 32–46.

92. Dallari, S., et al., Enteric viruses evoke broad host immune responses resembling those elicited by the bacterial microbiome. Cell Host Microbe, 2021. 29(6): p. 1014–1029.e8.

93. Friedman, E.S., et al., FXR-Dependent Modulation of the Human Small Intestinal Microbiome by the Bile Acid Derivative Obeticholic Acid. Gastroenterology, 2018. 155(6): p. 1741–1752.e5.

94. Wang, S., et al., Diet-induced remission in chronic enteropathy is associated with altered microbial community structure and synthesis of secondary bile acids. Microbiome, 2019. 7(1): p. 126.

